# Composition and Biophysical Properties of the Sorting Platform Pods in the *Shigella* Type III Secretion System

**DOI:** 10.1101/2021.05.13.444006

**Authors:** Shoichi Tachiyama, Ryan Skaar, Yunjie Chang, Brittany Carroll, Meenakumari Muthuramalingam, Sean K. Whittier, Michael L. Barta, Wendy L. Picking, Jun Liu, William D. Picking

## Abstract

*Shigella flexneri*, causative agent of bacillary dysentery (shigellosis), uses a type III secretion system (T3SS) as its primary virulence factor. The T3SS injectisome delivers effector proteins into host cells to promote entry and create an important intracellular niche. The injectisome’s cytoplasmic sorting platform (SP) is a critical assembly that contributes to substrate selection and energizing secretion. The SP consists of oligomeric Spa33 “pods” that associate with the basal body via MxiK and connect to the Spa47 ATPase via MxiN. The pods contain heterotrimers of Spa33 with one full-length copy associated with two copies of a C-terminal domain (Spa33^C^). The structure of Spa33^C^ is known, but the precise makeup and structure of the pods *in situ* remains elusive. We show here that recombinant wild-type Spa33 can be prepared as a heterotrimer that forms distinct stable complexes with MxiK and MxiN. In two-hybrid analyses, association of the Spa33 complex with these proteins occurs via the full-length Spa33 component. Furthermore, these complexes each have distinct biophysical properties. Based on these properties, new high-resolution cryo-electron tomography data and architectural similarities between the Spa33 and flagellar FliM-FliN complexes, we provide a preliminary model of the Spa33 heterotrimers within the SP pods. From these findings and evolving models of SP interfaces and dynamics in the *Yersinia* and *Salmonella* T3SS, we suggest a model for SP function in which two distinct complexes come together within the context of the SP to contribute to form the complete pod structures during the recruitment of T3SS secretion substrates.

## 1 Introduction

Type III secretion systems (T3SS) are used by many Gram-negative bacterial pathogens to inject host altering proteins into the cytoplasm of target cells (Portaliou et al., 2016; Galan and Waksman, 2018). The *Shigella flexneri* T3SS apparatus (injectisome) comprises four assemblies that work cooperatively to secrete effectors. The needle tip complex is exposed on the bacterial surface where it can make contact with the host cell membrane and contribute to the formation of a translocon pore (Deane et al., 2006; Espina et al., 2006b). The translocon and tip complex reside at the distal end of a needle that is generated from a polymer of MxiH (Cordes et al., 2003; Kenjale et al., 2005). The needle is then anchored to an envelope-spanning basal body (the injectisome core) (Blocker et al., 1999; Lunelli et al., 2020), which is associated with a cytoplasmic sorting platform (SP) (Hu et al., 2015). The SP contributes to substrate selection and to energizing secretion. The SP was originally proposed as a complex in *Salmonella* that included SctK, SctQ and SctL (Lara-Tejero et al., 2011) for which the *Shigella* homologues are MxiK, Spa33 and MxiN, respectively. An increasingly accepted uniform nomenclature for the injectisome components is provided as a guide in **Table 1** (Wagner and Diepold, 2020) and will be used here when referring to all non-*Shigella* T3SS.

**Table 1.** The standardized nomenclature for the inner membrane ring (IR) and sorting platform components for injectisome components discussed here.

| Organism | Outer<br>IR <sup>a</sup> | Inner<br>IR | Pod and<br>Adaptor<br>Proteins | Spokes | ATPase |
| --- | --- | --- | --- | --- | --- |
| **Standardized**<br>**Nomenclature <sup>b</sup>** | **SctD** | **SctJ** | **SctQ/SctK** | **SctL** | **SctN** |
| *Shigella* species | MxiG | MxiJ | Spa33/MxiK | MxiN | Spa47 |
| *Salmonella enterica* | PrgH | PrgK | SpaO/OrgA | OrgB | InvC |
| *P. aeruginosa* | PscD | PscJ | PscQ/PscK | PscL | PscN |
| *Yersinia* species | YscD | YscJ | YscQ/YscK | YscL | YscN |

<sup>a</sup> The outer portion of the inner membrane ring (IR) possesses a transmembrane helix that connects a periplasmic domain that assembles into a ring with 24-fold symmetry and a cytoplasmic portion that is a smaller forkhead associated-like (FHA-like) domain.
<sup>b</sup> A standardized nomenclature has been proposed using the common name Sct (secretion and cellular translocation) for each T3SA component (Wagner and Diepold, 2020).

A key interface between the basal body and the SP in *Shigella* is formed by interaction of the inner membrane ring (IR) protein MxiG (SctD) with an SP adaptor protein. The cytoplasmic forkhead associated-like (FHA-like) domain of MxiG (MxiG^C^) and the SP protein MxiK (SctK) mediate this interaction (Tachiyama et al., 2019). A feature of this interface is a switch from the 24-fold symmetry of the IR components to the six-fold symmetry of MxiK, which is in association with the SP “pods”. The pods are the largest densities in the SP and comprise heterooligomeric complexes of Spa33 (SctQ) (Hu et al., 2015; McDowell et al., 2016). Spa33, in turn, interacts with MxiN (SctL) and this links them to the central ATPase Spa47 (SctN) (**Suppl. Fig. S1**) (Hu et al., 2015; Notti et al., 2015). Our understanding of the composition of the SP has evolved with recent cryo-electron tomography (cryo-ET) work using minicells derived from *Shigella* (**Suppl. Fig. S1**), which are in agreement with structures seen using similar methods for the *Salmonella* injectisome (Hu et al., 2015; Hu et al., 2017). Interestingly, deletion of *mxiK* or *spa33* results in complete loss of SP assembly, however, mutants in which the radial spoke (and ATPase regulator) protein gene *mxiN* is deleted continue to allow remnants of the pods to be viewed while Spa47 is absent (**Suppl. Fig. S2**) (Hu et al., 2015).

Recombinant Spa33 (expressed from the wild-type gene) purifies as a heterotrimer comprising one copy of full-length Spa33 and two copies of a C-terminal fragment generated via an alternative internal translation start site (McDowell et al., 2016). The C-terminal fragment (Spa33^C^) is a “surface presentation of antigens” (SPOA) domain named for alternative forms of SctQ from *Salmonella* and in *Pseudomonas syringae* (Fadouloglou et al., 2004; Notti et al., 2015; Notti and Stebbins, 2016). As described in depth for SctQ from *Salmonella*, full-length Spa33 possesses two SPOA domains at its C-terminus, but an internal translation start site allows for the second of these (SPOA2) to be expressed separately from the full-length protein (Notti et al., 2015). SPOA domains have a significant β-structure component and are structurally equivalent to FliN from *Thermotoga maritima*, which is a component of the flagellar C-ring (**Suppl. Fig. S3**) (Brown et al., 2005; McDowell et al., 2016). Interestingly, the Spa33 heterotrimer shares significant similarity to the FliM-FliN heterotetramer (FliM-FliN_3_) where the full-length Spa33 contributes its C-terminal SPOA2 domain and the Spa33^C^ homodimer contributes two additional FliN-equivalent SPOA domains (Fadouloglou et al., 2004; McDowell et al., 2016). This observation prompted the Lea group to logically propose a model in which Spa33 forms a torus structure resembling the flagellar C-ring (McDowell et al., 2016). Subsequent cryo-ET imaging, however, indicated that the bulk of the SP is actually made up of discontinuous pods described above (Hu et al., 2015).

Diepold *et al*. estimated that there are approximately 22 copies of SctQ in the SP of *Yersinia* (Diepold et al., 2015). This is consistent with the observation that the crystal structure of Spa33^C^ dimers would only occupy a small portion of the pod densities attributed to Spa33. Based on available data, it now appears that the injectisome pods each consist of multiple copies of Spa33 and Spa33^C^. Furthermore, Diepold *et al*. found that during active secretion these SctQ components are in continual exchange between the injectisome SP and a distinct cytoplasmic pool of these proteins (Diepold et al., 2015; Diepold et al., 2017). The dynamics occurring within the SP ultrastructure are considered to be important for type III secretion system activity. Therefore, a better understanding of the structure of the SctQ complexes within the SP should move the field closer to understanding the mechanisms that govern injectisome function.

It was recently suggested that in *Salmonella*, the SctQ C-terminal domain (SctQ^C^) may not be required for generating a functional injectisome (Lara-Tejero et al., 2019). Instead, it may only stabilize full-length SctQ by occupying a site on the protein that within the functional SP is occupied by SctL. Bernal *et al*. suggested that a complex consisting of SctQ, SctQ^C^, SctL and SctN can be prepared, perhaps suggesting that SctQ^C^ is a required component of the fully functional injectisome (Bernal et al., 2019). Nevertheless, elimination of SctQ^C^ genetically still allowed for secretion of the *Salmonella* invasion proteins (Sips) and only reduced invasiveness by 50%, which is in agreement with the work suggesting SctQ^C^ is not essential. On the other hand, Bernal *et al*. did report that elimination of the internal translation start site still allowed for low-level production of SctQ^C^ (Bernal et al., 2019). Work in the *Shigella* system suggests that Spa33^C^ is indeed required for forming a functional SP because a mutation that eliminates the internal alternative translation start site to limit expression to full-length Spa33 (Spa33^FL^) fails to restore detectable secretion of translocator and effector proteins (McDowell et al., 2016). The contrasting findings indicate a need to better understand the actual makeup and organization of the SP pods in *Shigella*. Even more recently, additional Spa33 protein variants were identified, including a 6.5 kDa form called Spa33^CC^ (found to be important for injectisome activity) and two other Spa33 variants (called Spa33^N^ and Spa33^X^) that have no obvious functions and may be translational artifacts (Kadari et al., 2019). Although we have also observed production of Spa33^CC^ (unpublished observation), it will not be considered further here since it does not associate with the Spa33 heterotrimer.

While the *Shigella* and *Salmonella in situ* injectisomes are similar in structure based on cryo-ET imaging (Hu et al., 2015; Hu et al., 2017), there are differences between the two that warrant investigating both. The functional *Shigella* injectisome requires expression of wild-type *spa33* and the production of one Spa33^FL^ and two copies of Spa33^C^ (McDowell et al., 2016). Moreover, we show here that the Spa33 heterotrimer forms stable complexes with MxiK and MxiN that have distinct biophysical behaviors *in vitro*. Within these complexes, Spa33^C^ acts to stabilize Spa33^FL^, which is otherwise not soluble, and the Spa33 heterotrimer stabilizes MxiK, which also is not otherwise soluble. Two-hybrid analyses indicate that Spa33^FL^ is able to associate with MxiK, but Spa33^C^ is not, suggesting that Spa33^FL^ mediates formation of the Spa33-MxiK complex. Meanwhile, it was wild-type Spa33 that was found to purify as part of the complex with MxiK, suggesting that Spa33^C^ is required to stabilize this overall complex, as well. Likewise, MxiN specifically interacted with Spa33^FL^ according to two-hybrid analyses but not with Spa33^C^, and it could associate with the Spa33 heterotrimer to form Spa33-MxiN complexes *in vitro*. Based on our ability to generate these complexes, their biophysical properties and old and new higher-resolution cryo-ET images, we propose that these two Spa33-containing complexes have critical discrete roles in active type III secretion.

## 2 Materials and Methods

### 2.1. Materials and strains

*S. flexneri* 5a (M90T) was provided from J. Rohde (Dalhousie University, Halifax, NC, Canada). *Shigella mxiK* and *mxiN* null strains were from A. Allaoui (Université Libre de Bruxelles, Bruxelles, Belgium) the and *spa33* null strain was from C. Lesser (Massachusetts General Hospital, Cambridge, Massachusetts, USA). Mutant strains were grown at 37 °C on trypticase soy agar (TSA) plates containing Congo red and then used to inoculate trypticase soy broth (TSB) containing 50 µg/mL kanamycin to select for virulence plasmid-harboring null strains. Ampicillin (100 µg/mL) was added to select wild-type *mxiK*, *mxiN*, *spa33*, or their mutant genes in the pWPsf4 expression plasmid. To generate minicells, wild-type *S. flexneri* was transformed with pBS58, which is a low-copy number plasmid encoding the spectinomycin-resistance gene. pBS58 continuously expresses the *E. coli* genes *ftsQ*, *ftsA*, and *ftsZ*, which causes abnormal cell division to produce *Shigella* minicells (Bi and Lutkenhaus, 1990; Hu et al., 2015; Farley et al., 2016). To produce *Shigella* minicells, spectinomycin (100 µg/mL) was added into trypticase soy broth (TSB) to grow the strain for overnight. A mL of this overnight culture was added into 100 mL of fresh TSB with spectinomycin (100 µg/mL) and grown at 37 °C to an absorbance of 1.0 at 600 nm (A_600_) (Hu et al., 2015; Tachiyama et al., 2019). To enrich for minicells, the culture was centrifuged at 1000 × g for 5 min to remove the large cells, and the supernatant fraction was further centrifuged at 20,000 × g for 10 min to collect the minicells for subsequent cryo-ET imaging of the SP pods.

### 2.2. Protein expression

To prepare expression vectors with wild-type *spa33*, *spa33^C^*, *spa33^FL^* (lacking the internal translation start site), *mxiK* or *mxiN*, each gene from *S. flexneri* was amplified using PCR. PCR products were purified and inserted into the pT7HMT expression vector (Geisbrecht et al., 2006; Tachiyama et al., 2019), which encodes His_6_-tag with TEV protease cleavage site at N-terminal side of the cloning site and a kanamycin resistance gene. To generate *E. coli* strains for protein expression, these recombinant plasmids were introduced into Tuner (DE3) cells (Novagen). For co-expression of *mxiK* and *spa33*, the latter was inserted into multiple cloning site two in pACYCDuet^TM^ (Novagen). *E. coli* Tuner(DE3) cells were then transformed with both *mxiK* in pT7HMT and *spa33* in pACYCDuet^TM^. To induce protein expression, *E. coli* strains with recombinant pT7HMT vector were inoculated into LB with kanamycin (50 µg/mL) and grown overnight. The overnight bacterial culture was used to inoculate a larger volume of LB with the antibiotic. When the A_600_ reached 0.6, the cultures were induced with 0.5 mM isopropyl β-D-1-thiogalactopyranoside (IPTG). The culture was then incubated at 16 °C overnight. MxiK and Spa33 production was carried out the same way except that Terrific Broth (TB) with kanamycin and chloramphenicol (34 µg/mL) was used for the large volume culture with induction started at an A_600_ of 1.0.

### 2.3. Protein Purification

The cells expressing the protein of interest were harvested by centrifugation at 4000 rpm for 10 min at 4 °C. The cell pellet was suspended in immobilized metal affinity chromatography (IMAC) binding buffer (5 mM imidazole, 0.5 M NaCl, 20 mM Tris) pH 7.5. The lysate was prepared using a microfluidizer at 15,000 psi for three cycles and clarified by centrifugation at 10,000 rpm for 30 min at 4 °C. The clarified supernatant was applied to a Ni^2+^-charged IMAC column with chromatography carried out on an AKTA^TM^ purification system (GE Healthcare, Chicago, IL). The loaded column was washed with additional IMAC binding buffer and then elution carried out using an elution buffer (0.5 M imidazole, 0.5 M NaCl, and 20mM Tris, pH 7.5). The His_6_ tag was removed using TEV protease after overnight dialysis in IMAC binding buffer. The protein was then applied to a second Ni-IMAC column and the flow-through collected, concentrated and buffer exchanged into 20 mM Tris, 200 mM NaCl, pH 7.5 prior to loading onto an equilibrated Superdex 200 pg (GE) for size exclusion chromatography (SEC). All fractions were analyzed by SDS-PAGE. SEC was also used to purify Spa33-MxiN complexes after mixing the proteins after each had been purified separately. TheMxiN, Spa33-MxiK and Spa33-MxiN samples were made to 10% (v/v) glycerol.

### 2.4. Contact-mediated hemolysis

To test for secretion and translocon formation, contact-mediated hemolysis was performed for each *Shigella* mutant strain. Each strain was grown on a trypticase soy agar (TSA) plate containing Congo-red (Picking et al., 2005; Tachiyama et al., 2019). Isolated colonies were inoculated into TSB for growth at 37 °C until mid-log phase with the appropriate antibiotics added for selection. A 10 mL aliquot of each culture was centrifuged at 4,000 rpm for 10 min at 30 °C and the pellet was resuspended in 10 mM phosphate, pH 7.4 containing 150 mM NaCl (PBS). The PBS was adjusted to give equal concentrations of bacteria in each case based on the A_600_. To prepare red blood cells, defibrinated sheep red blood cells (Colorado Serum, Co.) were washed with PBS and resuspended into 3 mL PBS. Then 50 µL of the resuspended bacteria were added into 50 µL of washed red blood cells in a 96-well plate. The plate was centrifuged at 3,500 rpm for 15 min at 30 °C to force *Shigella* contact with red blood cells. The plate was then incubated at 37 °C for an hour. The cell mixtures were then vigorously resuspended with 100 µL cold PBS and centrifuged at 3,500 rpm for 15 min at 10 °C. The supernatants were then transferred into a new 96-well plate and the amount of released hemoglobin was measured based on the A_545_. The absorbance was used to calculate relative hemolysis for each strain after comparison to a control for negative hemolysis (incubation in PBS) and a control for positive (complete) hemolysis (incubation in water).

### 2.5. Far-UV circular dichroism (CD) spectroscopy

The secondary structure of recombinant proteins and their complexes were analyzed by CD spectroscopy (Espina et al., 2006a; Birket et al., 2007) using a JASCO model J-1500 CD spectropolarimeter (Jasco Inc., Easton, MD). Samples were prepared in 20 mM Tris, pH 7.0, 200 mM NaCl, 5.0 % glycerol and the same buffer was used for blank subtraction. For MxiN, tris(2-carboxyethyl)phosphine (TCEP) was added to the buffer to a final concentration of 1 mM. For Spa33 and Spa33^C^, PBS was used as the buffer. To obtain CD spectra, ellipticities between 190 and 260 nm from each protein sample in 0.1 cm path length quartz cuvettes were obtained at 10 °C. Molar ellipticities were then calculated based on protein concentration, molecular weight, and residue numbers. To monitor thermal stability, the molar ellipticity at 222 nm (a minimum seen for proteins having an α-helical component) was monitored every 2.5 °C with increasing temperature from 10 to 90 °C at a ramp rate of 15 °C/h rate. The molar ellipticity was then plotted as a function of temperature to provide a thermal unfolding curve. Thermal transition points or unfolding midpoints (Tm) was determined using Prism software from GraphPad.

### 2.6. Cryo-ET data collection and tomogram reconstruction

A Polara G2 electron microscope (FEI Company) equipped with a field emission gun and a Direct Detection Camera (K2 summit, Gatan) was used to image frozen-hydrate minicells from the wild-type *Shigella* strain at −170 °C. The microscope was operated at 300 kV, with a pixel size of 2.6 Å at the specimen level. The single-axis tilt series was collected at about 5 µm defocus using SerialEM software (Mastronarde, 2005) and with a cumulative dose of ∼50 e/Å^2^ distributed over 35 stacks covering an angle range of −51° to 51° with 3° angle step.

Motioncorr2 (Zheng et al., 2017) was used to do drift correction for each image stack. Then, the tilt series were aligned by IMOD (Kremer et al., 1996), and weighted back-projection (WBP) or Simultaneous Iterative Reconstruction Technique (SIRT) was used to reconstruct the tomograms by Tomo3D software (Agulleiro and Fernandez, 2015). A total of 922 tomograms were reconstructed for the wild-type *Shigella* minicell sample.

### 2.7. Sub-tomogram averaging

A total of 4,488 injectisome particles from wild-type *Shigella* were manually picked from the SIRT reconstructed tomograms as previously described (Hu et al., 2015). The particle positions were then used to extract the sub-tomograms from the 1×1×1 binned WBP reconstructed tomograms. I3 software (Winkler, 2007; Winkler et al., 2009) was used for the sub-tomogram averaging. 4×4×4 binned sub-tomograms were used for an initial alignment and 3D classification, and then 2×2×2 (or 1×1×1) binned sub-tomograms were used for refinement. To improve the resolution of the pod structure, a single pod region in the subtomogram average (the green arrow region in Fig. 7 A) was extracted for additional focused refinement processes.

### 2.8. 3D visualization and molecular modeling

The SP of the overall T3SA (see **Fig. 7A**) and the refined single pod structures were used to create 3D surface rendering images of the sub-tomogram averaging using UCSF Chimera (Pettersen et al., 2004) and ChimeraX (Goddard et al., 2018) software. For the molecular model, the crystal structure of SctK from *P. aeruginosa* (Muthuramalingam et al., 2020) was fitted into the MxiK position in the refined pod rendering image using “fit in map” in UCSF Chimera (Pettersen et al., 2004; Hu et al., 2015; Tachiyama et al., 2019; Muthuramalingam et al., 2020). Three FliN (PDB: 1YAB) (**Suppl. Fig. S3**) and a FliM (PDB: 2HP7) (see **Suppl. Fig. S14**) crystal structures from *Thermotoga maritima* were used to generate a Spa33 heterotrimer model based on a FliN and FliM complex model from the flagellar C-ring (Carroll et al., 2020). Two copies of this model could fit into the bottom part of Spa33 region using the “fit in map’ in UCSF Chimera software (Pettersen et al., 2004). The single pod surface rendering image was then fit back into six pod regions in the 3D surface rendering image of the SP (see **Fig. 7**).

### 2.9. Data and software accessibility

The pod cage structure and the refined images have been deposited in the EM Data Bank (EMD-23440).

### 2.10. Bacterial adenylate cyclase two-hybrid (BACTH) assay

A BACTH assay was used to assess possible interactions between the different SP proteins (MxiK, MxiN and Spa33 and their mutants) in an *in vivo*-like environment (Tachiyama et al., 2019). CyaA subdomain (T25 and T18) fusion proteins were prepared by cloning the proteins of interest from wild-type bacteria. The subdomains were fused to either the N- or C-terminus of the potential interacting protein. The pKT25 expression vector fuses the T25 subdomain to the protein’s N-terminus while the pKNT25 vector fuses the T25 subdomain to the C-terminal end of the protein. The same is achieved for fusion of the T18 subunit using the pUT18 or pUT18C vectors, which fuses the T18 unit to the N- and C-terminus of the protein.

Two compatible plasmids containing the gene fusions of interest were used to co-transform the *cyaA*^-^ *E. coli* strain, BTH101. BTH101 was co-transformed with empty plasmids was used for the negative control and the strain was co-transformed with plasmids encoding leucine zipper binding partners for the positive control. Individual colonies were selected on LB plates containing ampicillin and kanamycin and these were inoculated into LB broth containing these antibiotics and 0.5 mM IPTG. The bacteria were grown at 30 °C overnight. The overnight bacterial cultures were washed three times with M63 minimum medium and 2 µL was spotted onto indicator plates. Indicator plates could be: a) LB agar plates with ampicillin, kanamycin, 40 µg/mL X-Gal and 0.5 mM IPTG; b) MacConkey agar plates with ampicillin, kanamycin and 1.0 % (w/v) maltose; or c) M63 plates with ampicillin, kanamycin, 40 µg/mL X-Gal and 0.5 mM IPTG. These plates were incubated at 30 °C for four to five days and color changes over time were compared with that seen for the controls.

### 2.11. Testing the MxiN-Spa33 interaction using size-exclusion chromatography (SEC)

Purified MxiN and Spa33 were both dialyzed into 20 mM Tris, pH 7.5, 150 mM NaCl with 10% glycerol (v/v) for SEC. The MxiN and Spa33 were combined with an excess of MxiN and the mixture was concentrated to a final volume of 2 mL for injection onto a Hi Load 26/600 Superdex 200 pg column (GE Healthcare Life Sciences). The protein absorbance at 280 nm was monitored to detect protein elution on an AKTA system (GE Healthcare Life Sciences). The eluted fractions were collected for SDS-PAGE analysis.

### 2.12. Biolayer Interferometry (BLI)

The binding of MxiN to the wild-type Spa33 was assessed by biolayer interferometry (BLI) as described elsewhere (Barta et al., 2017). Purified His_6_-tagged MxiN was dialyzed into PBS with 10% glycerol, 0.01% BSA and 0.002% Tween 20 (BLI buffer). The MxiN was then immobilized onto a Ni-NTA biosensor tip and washed with BLI buffer for 5 min to remove non-immobilized protein. Once a baseline signal was established, the sensors with immobilized protein were immersed into a solution containing untagged Spa33, which was prepared and adjusted to 400, 200, 100, 50, 25 and 12.5 µg/mL. The Octet RED96 system (FortéBIO, Inc., Freemont, CA) was then used to monitor the binding of Spa33 to MxiN that had been immobilized onto a Ni-NTA biosensor tip. The binding of Spa33 at each concentration to the immobilized MxiN was measured in real time to obtain the association rate. The biosensor tips were then placed into BLI buffer lacking Spa33 to measure the dissociation rate. All steps were performed at 25 °C. Real-time data were analyzed using Octet Software version 8.2 (ForteBio). Binding kinetics (association and dissociation) as well as steady-state equilibrium concentrations were fitted using a 1:1 Langmuir binding model. Background correction was done by subtracting the data from ligand-loaded biosensor that had been incubated without analyte (Spa33). All steps were carried out with constant shaking at 1000 rpm and each measurement was performed in triplicate with an association time of 300 s followed by 300 s dissociation. The sensorgrams from the experiment were fitted with 1:1 binding equations available for interaction from a global curve fit using Data Analysis software 7.1.0.36 (ForteBio) (Barta et al., 2017).

### 2.13. Dynamic Light Scattering (DLS)

The effect that Spa33 inclusion into complexes with MxiK and MxiN has on particle size and surface charge was determined using dynamic light scattering (DLS). DLS measurements were made on a Malvern ZetaSizer Ultra using a quartz cuvette with a 1-cm path length. Backscattering was detected at an angle of 174.7 degrees, with each measurement performed in triplicate. Size distributions were calculated using the ZS Xplorer software (Malvern). The same instrument was used to determine the zeta-potential for each protein and protein complex.

## 3 Results

### 3.1. Interactions within the Spa33 heterotrimer

When Spa33 is expressed in *E. coli*, it readily purifies as a complex containing one full-length Spa33 (Spa33^FL^) with two copies of Spa33^C^, which are expressed from an internal translation initiation site (**Suppl. Fig. S4**). When the alternative start site is mutated to prevent initiation of the internal translation event, giving only Spa33^FL^, the protein could not be stably expressed and purified from *E. coli*, due to the fact that it is insoluble and sequestered to inclusion bodies. This is in contrast to expressing a gene that allowed for expression of only Spa33^C^, which is then easily purified as an apparent homodimer (**Suppl. Fig. S4**). Spa33^C^ was previously used to obtain a crystal structure for this SPOA domain (McDowell et al., 2016). The fact that Spa33 forms an oligomer is further supported by data from a bacterial adenylate cyclase two-hybrid (BACTH) assay where both the bait and prey plasmids contained the wild-type *spa33* gene (**Table 2** and **Suppl. Fig. S5, top**). The results with wild-type Spa33 are somewhat complicated since when the two adenylate cyclase domains are placed at the Sp33 N-terminus an interaction was observed. This could suggest that multiple Spa33 complexes were forming a higher order complex. We thus felt that it was imperative to look at interactions involving only Spa33^FL^, only Spa33^C^, and Spa33^FL^ plus Spa33^C^.

**Table 2.**
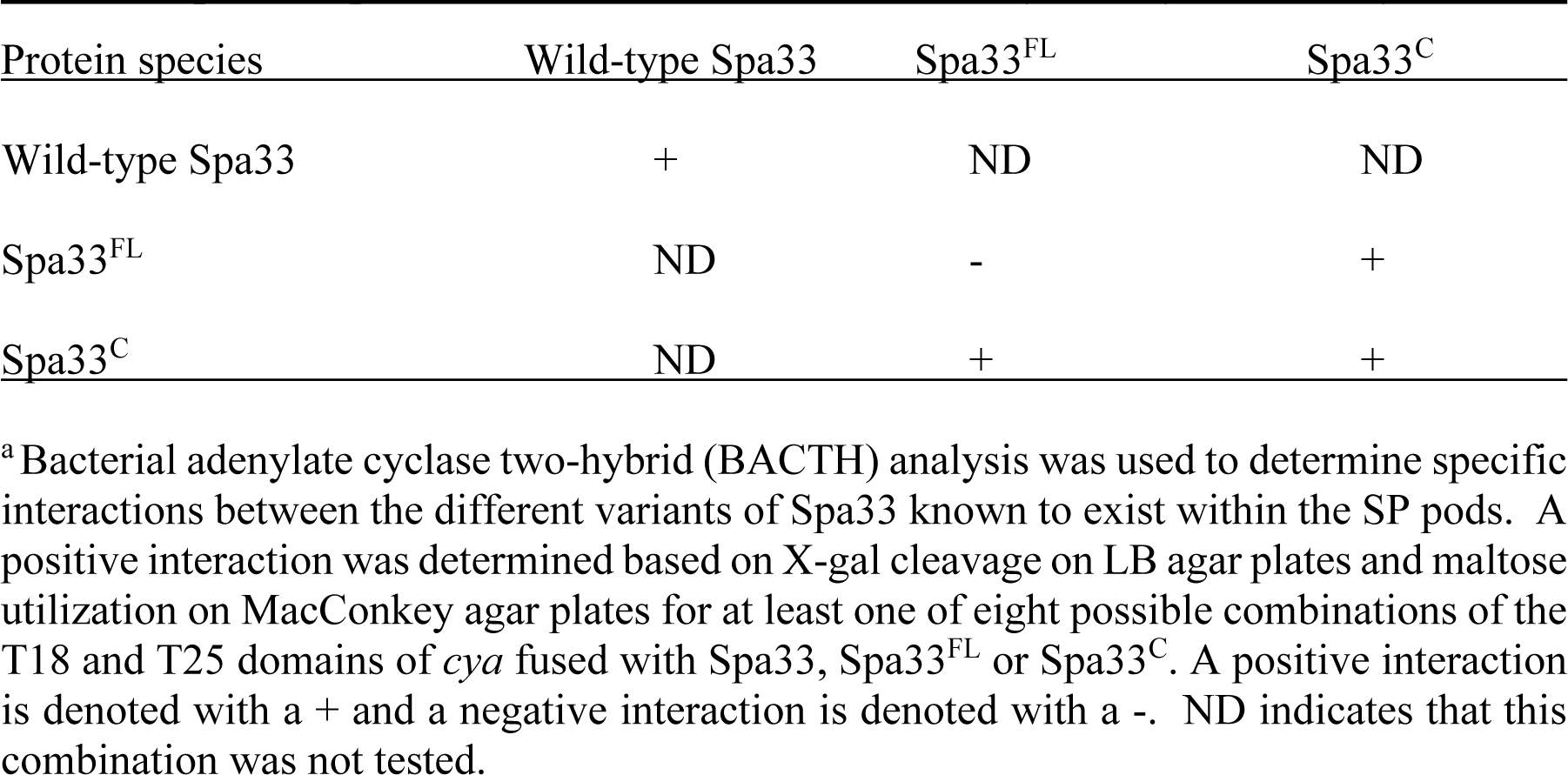
Specific Spa33 interactions can be identified by two-hybrid^a^ analysis.

When Spa33^FL^ was used in the same two-hybrid assay, no interaction was detected for any of the eight fusion permutations with the T18 and T25 fragments of the adenylate cyclase (**Table 2** and **Suppl. Fig. S5, bottom**). In contrast, when Spa33^C^ was used in the same assay, a clear interaction was again observed (**Table 2** and **Suppl. Fig. S6**). It is worth noting that not every fusion permutation between the Spa33^C^ binding pairs gave a positive reaction, but this is not surprising given that the interaction between the binding partners must orient the T18 and T25 fragments of the CyaA adenylate cyclase domains properly for activity to be restored (**Suppl. Fig. S7**). Perhaps more surprising was the observation that all combinations of wild-type Spa33 were positive for interactions in the BACTH system (**Suppl. Fig. S5**). Finally, when Spa33^FL^ was used in the same two-hybrid analysis with Spa33^C^, an interaction was observed, though not for every fusion permutation, which is consistent for what was seen for the Spa33^C^-Spa33^C^ interaction (**Table 2** and **Suppl. Fig. S8**). These data confirm what is currently believed to occur with respect to the interactions within the Spa33 heterotrimer. However, they also show that interactions within the Spa33-containing pods of the SP do not arise from Spa33^FL^-Spa33^FL^ interactions, at least not in the absence of Spa33^C^.

It was previously reported that neither Spa33^FL^ nor Spa33^C^ alone could restore the secretion of Ipa proteins when expressed in a *Shigella spa33* null strain (McDowell et al., 2016). In agreement with this prior observation, neither of these forms of Spa33 alone could support contact-mediated hemolysis, although a small amount of residual hemolytic activity was observed for Spa33^FL^ (**Fig. 1**). The reason for the low level of hemolysis for Spa33^FL^ was not entirely clear. It is possible that it could be related to the reported observation by the Lara-Tejero *et al*. that the Spa33^C^ homologue from *Salmonella* (SctQ^C^) is not required for secretion of translocator proteins or for cellular invasion (Lara-Tejero et al., 2019) and that it merely contributes to stabilization of SctQ^FL^ (the full-length form). They also suggested that SctQ^C^ contributed to the stabilization of other SP components (SctK and SctL). Indeed, Spa33^C^ is required for stabilization of Spa33^FL^ since in its absence Spa33^FL^ is insoluble, however, the very minor degree of restoration of contact-mediated hemolysis for *Shigella* expressing Spa33^FL^ is not at a level considered here to indicate it gives rise to an active injectisome. This is consistent with our observation that when the contact-hemolysis assay is allowed to incubate for long periods of time, the level of hemoglobin release gradually increases (unpublished observation). Thus, there may be degradation of components (*i.e.* Spa33^FL^) that allow for undetectable amounts of Spa33^C^ to accumulate and allow residual T3SS activity to be seen as a function of time. This would fit with observations by Bernal *et al*. who found that SctQ^FL^ gave rise to a minor population of SctQ^C^ (Bernal et al., 2019), however, even in that report there was a 50% restoration of injectisome function by the SctQ^FL^ and we see nothing near that level of contact-mediated hemolysis for Spa33^FL^ (**Fig. 1**).

**Figure 1.**
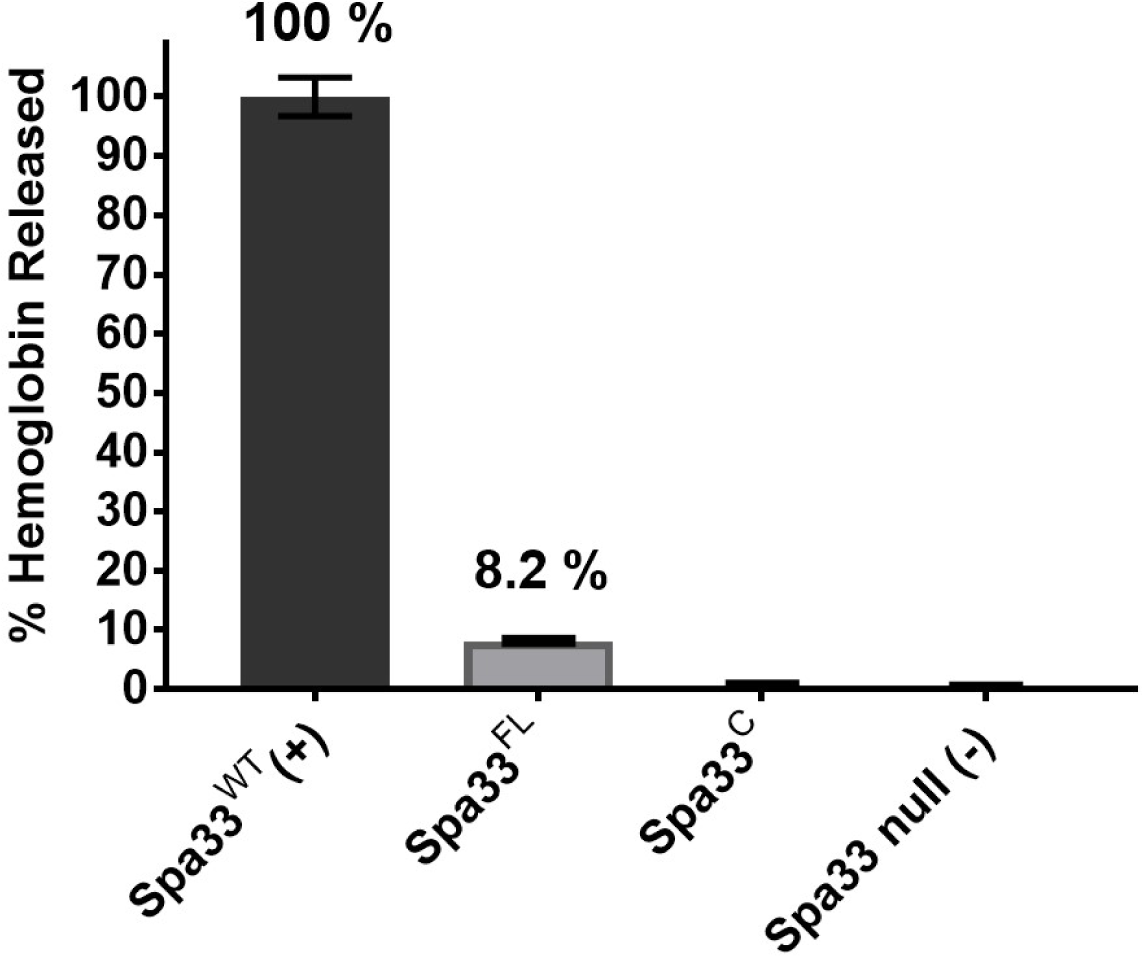
Contact-mediated hemolysis shows that Spa33^FL^ and Spa33^C^ are not active when expressed in a *Shigella spa33* null mutant. To test the type III secretion activities of the Spa33 variants *in vivo*, contact-mediated hemolysis was used to measure of translocon formation. Spa33^FL^ did restore a minor amount of hemolysis activity (about 8%), however, Spa33^C^ was completely inactive. The level of injectisome activity seen for Spa33^FL^ was far below that seen for SctQ^FL^ in *Salmonella*. Statistical differences between wild-type Spa33 (Spa33^WT^) and other samples (in triplicate) were assessed using one-way ANOVA with Spa33^FL^, Spa33^C^ and the *spa33* null all with p <0.005) and error bars depict mean ± SD.

### 3.2. Biophysical analysis of wild-type Spa33 and Spa33^C^ using circular dichroism (CD) spectroscopy

Circular dichroism (CD) spectroscopy was used to compare the secondary structure properties of the purified Spa33 heterotrimer and Spa33^C^ homodimer. Both complexes possess a dominant β-structure component with only a minor amount of α-helical structure in each (**Fig. 2**). By monitoring the CD signal 222 nm as a function of temperature, we found that Spa33^C^ was highly stable with an unfolding transition midpoint (Tm) of 65 °C (**Fig. 2B** and **Table 3**). Meanwhile, the wild-type Spa33 had a Tm of 62 °C. Because Spa33^FL^ could not be purified as a recombinant protein, Spa33^C^ clearly had a significant solubilizing effect on it. In this respect, Spa33^C^ has a similar effect on Spa33^FL^ stability as does SctQ^C^ on SctQ^FL^ in *Salmonella* (Lara-Tejero et al., 2019).

**Figure 2.**
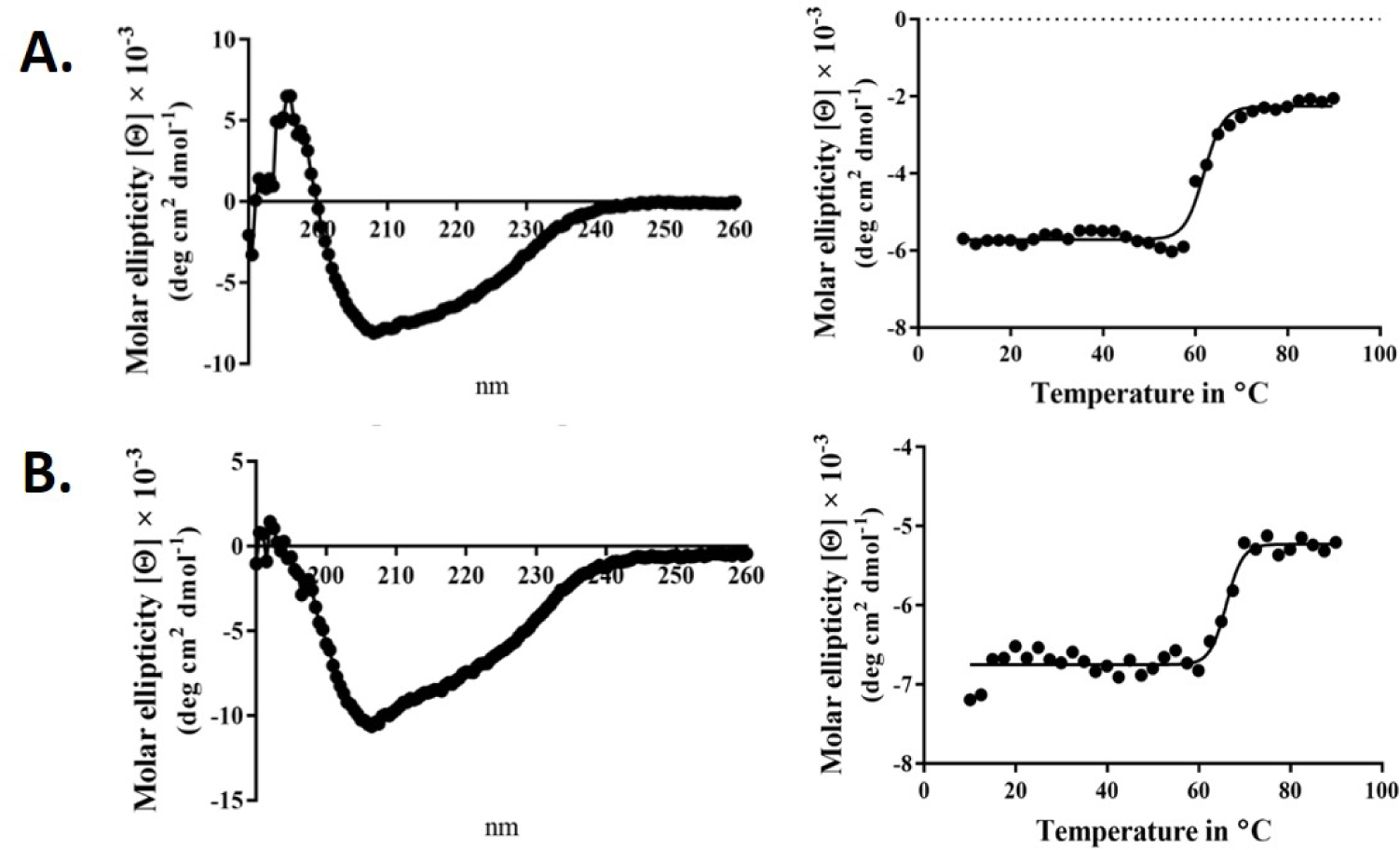
Circular dichroism (CD) spectra and thermal unfolding for wild-type Spa33 and Spa33^C^. CD spectroscopy was used to analyze the secondary structure content and thermal stabilities of wild-type Spa33 and Spa33^C^. **A**) The CD spectrum of wild-type Spa33 shows a negative peak at 208 nm with a modest shoulder at 222 nm (**left**). This spectrum is indicative of a significant amount of β-structure. To determine the Tm for wild-type Spa33, the molar ellipticity at 222 nm was monitored as a function of temperature. Based on the large transition observed (**right**) the Tm was found to be 61.88 °C. **B**) The CD spectrum of Spa33^C^ also shows more dominant 208 nm minimum suggesting a dominant β-structure component (**left**). Based on the thermal unfolding curve for Spa33^C^ (**right**) the Tm was determined to be 65.12 °C.

**Table 3.** Biophysical properties of Spa33, Spa33-containing complexes and MxiN.

| Protein or protein complex | T<sub>m</sub> (°C) <sup>a</sup> | Hydrodynamic diameter (nm) <sup>b</sup> | Zeta potential (mV) |
| --- | --- | --- | --- |
| Spa33 (wild-type) | 61.88 | 13.4 ± 0.43 | -8.87 ± 1.09 |
| Spa33<sup>C</sup> | 65.12 | 6.09 ± 0.11 | -6.16 ± 1.44 |
| Spa33/MxiK | 52.46 | 10.2 ± 0.66 | -7.25 ± 0.21 |
| MxiN | 36.63 | 9.85 ± 0.38 | -0.28 ± 0.20 |
| Spa33/MxiN | 38.99 | 18.5 ± 0.96 | -4.95 ± 0.82 |

<sup>a</sup> Unfolding temperature (T<sub>m</sub>) was determined as the midpoint of the thermal transition seen at 222nm using CD spectroscopy. In all cases, the R<sup>2</sup> was >0.98.
<sup>b</sup> Dynamic light scattering and zeta-potential measurements were carried out on a Zetasizer Nano (Malvern Instruments) at room temperature in PBS.

### 3.3. The formation of a Spa33 complex with MxiK

We have shown that MxiK interacts with wild-type Spa33 in the *in situ* injectisome and this was confirmed by two-hybrid analysis (Tachiyama et al., 2019). As a follow up, we carried out BACTH analyses using MxiK with wild-type Spa33, Spa33^FL^ and Spa33^C^. While we know that MxiK binds to wild-type Spa33, it has not been shown whether this interaction occurs through Spa33^FL^ or Spa33^C^ (summarized in **Table 4**). Based on two-hybrid analysis, we found that MxiK was able to interact with Spa33^FL^, however, we could not detect any interactions with Spa33^C^ (**Suppl. Fig. S9**). Whether this interaction involves the Spa33 N-terminal domain or the SPOA1-SPOA2 domains of Spa33^FL^ is not yet clear. In parallel experiments, we co-expressed wild-type *spa33* and *mxiK* in *E. coli* and found that the Spa33 heterotrimer could be co-purified as a complex with MxiK possessing an N-terminal His_6_-tag using IMAC followed by size-exclusion chromatography (**Fig. 3** and **Suppl. Fig. S10**). The fact that MxiK is part of this complex while it cannot be stably purified as a soluble protein on its own suggests a stabilizing, or minimally a solubilizing, effect for the Spa33 heterotrimer on MxiK.

**Figure 3.**
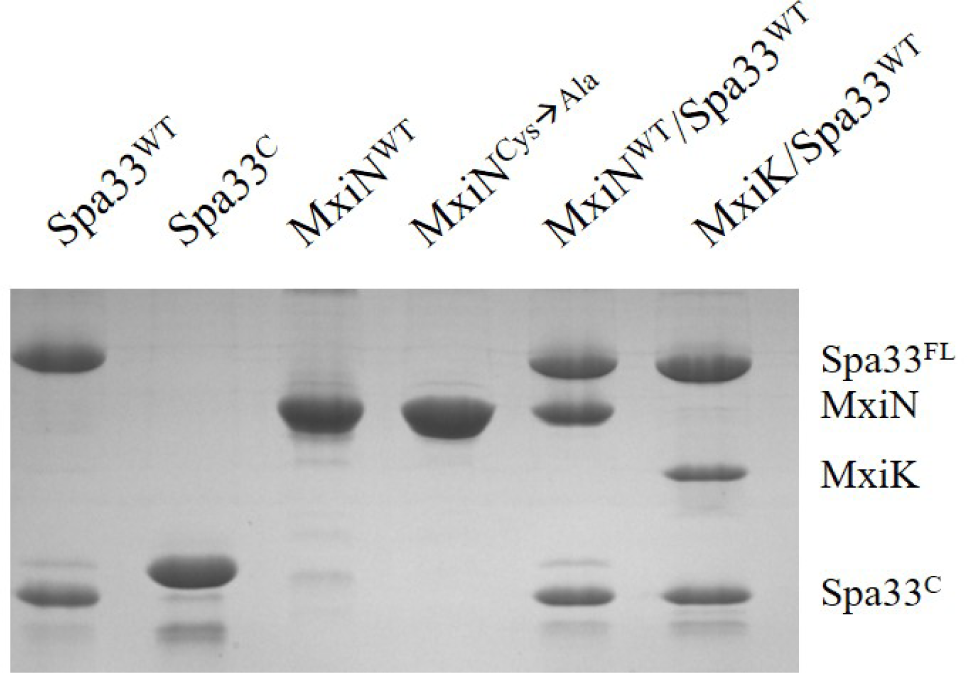
SDS-PAGE analysis for all purified complexes containing Spa33. From the left, wild-type Spa33, Spa33^C^, MxiN, a MxiN mutant with all Cys residues removed to allow future crystallography screens (fully active *in vivo*), the MxiN-Spa33 complex formed *in vitro* and the MxiK-Spa33 complex purified after co-expression in *E. coli.* The Spa33^C^ (second lane) is slightly larger than the same component in the other lanes because it still possesses an N-terminal His_6_-tag. Recombinant Spa33^FL^ and MxiK alone are not soluble and could not be purified.

**Table 4.** Interactions of MixK and MxiN with Spa33 and the individual components of the Spa33 complex.

| Protein 1 | Protein 2 | BACTH Result <sup>a</sup> |
| --- | --- | --- |
| Wild-type Spa33 | MxiK | + |
| Spa33<sup>FL</sup> | MxiK | + |
| Spa33<sup>C</sup> | MxiK | - |
| Wild-type Spa33 | MxiN | + |
| Spa33<sup>FL</sup> | MxiN | + |
| Spa33<sup>C</sup> | MxiN | - |
| MxiK | MxiK | - |
| MxiN | MxiN | + |

<sup>a</sup> Bacterial adenylate cyclase two-hybrid (BACTH) analysis was used to determine specific interactions between the Spa33 components known to exist within the SP pods and MxiN and MxiK as determined by BACTH analyses and shown in the **Supplemental Materials**. As in **Table 2**, a positive reaction for at least one of eight possible combinations of the T18 and T25 domains was considered to give a positive interaction for the pair. MxiK and MxiN were also tested for their ability to interact with themselves.

When the CD spectrum of the Spa33-MxiK complex was obtained, it indicated the presence of both α-helical and β-strand components. This is consistent with contributions made to secondary structure by both components of the complex (**Fig. 4**). We know that Spa33 is rich in β-structure and the crystal structure and CD spectrum of the MxiK homologue SctK from *P. aeruginosa* show that it is rich in α-helical structure (Muthuramalingam et al., 2020). When the signal at 222 nm was followed as a function of temperature, the complex showed a single unfolding transition at 52 °C though there may be a second minor transition occurring at about 60 °C (**Fig. 4** and **Table 3**). The main transition point or Tm is below that observed for the Spa33 heterotrimer, presumably because of structural rearrangements in Spa33 that occur upon complex formation that causes the complex to behave as a single entity with regard to thermal stability. This said, it can’t be ruled out that the minor second transition could be due to some of the Spa33 heterotrimer or Spa33^C^ homodimer escaping in a folded state during the first transition to later unfold at its characteristic Tm. The relative size of this second transition, however, is consistent with this only represented a small subpopulation of the Spa33 present in the original Spa33-MxiK complex. Therefore, because a single dominant Tm is observed and MxiK cannot be prepared as a recombinant protein by itself, the data support the proposal that the Spa33-MxiK complex mostly behaves as a single unit consisting of Spa33^FL^, Spa33^C^ and MxiK. Although not directly applicable here, MxiK can be purified in small amounts with a C-terminal T4L genetically fused to it (MxiK-T4L-C) and this fusion retains activity when expressed in a *Shigella mxiK* null mutant (Tachiyama et al., 2019). This purified chimera has a Tm of about 39 °C, which compares well with the Tm of SctK *P. aeruginosa*, which is 40 °C (Muthuramalingam et al., 2020). Thus, although the complex has a thermal unfolding transition below that of the Spa33 heterotrimer, it does have a significant stabilizing effect on MxiK. Likewise, the fact that a second transition is not observed supports the idea that it behaves as a single stably folded unit.

**Figure 4.**
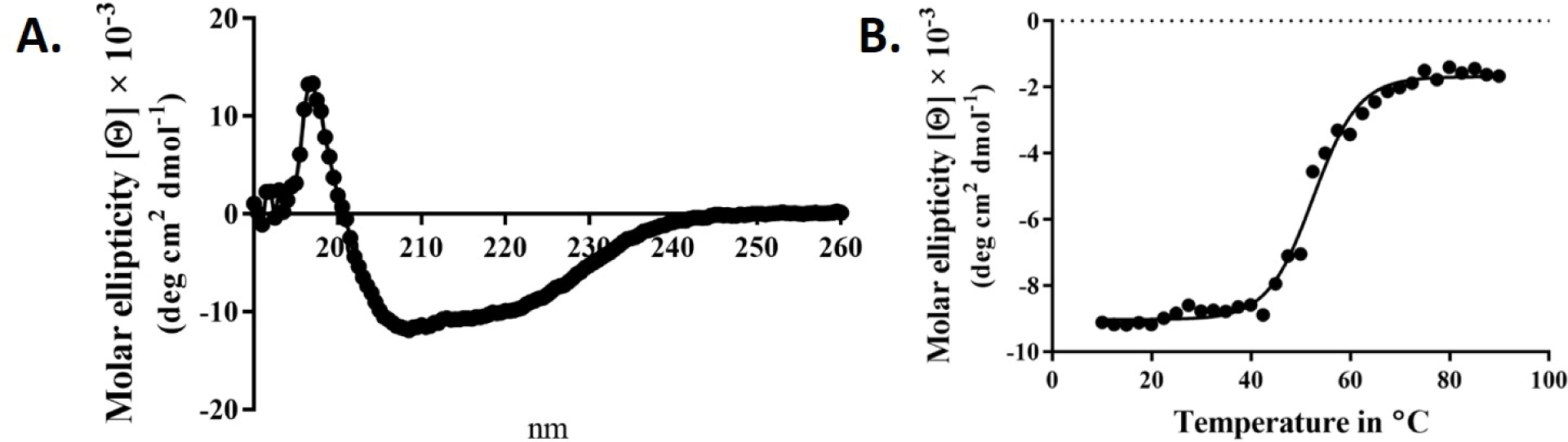
CD spectrum of the wild-type Spa33-MxiK complex. **A**) The secondary structure of the Spa33-MxiK complex shows a greater α-helical character than the Spa33 heterotrimer alone. **B**) To determine thermal stability, molar ellipticity at 222 nm was monitored as a function of temperature. Based on the large thermal transition observed, the complex had a Tm of approximately 52.46 °C. This Tm is lower than that of wild-type Spa33 (62 ⁰C) and higher than that of the chimera MxiK-T4L-C (39 °C), suggesting that the Spa33 interaction enhances MxiK solubility and thermal stability. R^2^ = 0.99.

### 3.4. Assembly of a Spa33 complex with MxiN *in vitro*

MxiN forms the radial spokes that connect the Spa33 pods of the SP with the hub which is a hexamer of the Spa47 ATPase. MxiN forms a dimer and interacts with wild-type Spa33 as observed in two-hybrid analyses (Tachiyama et al., 2019). When the same two-hybrid system was used to explore MxiN interactions with the two components of the Spa33 heterotrimer (summarized in **Table 4**), MxiN was found to interact with Spa33^FL^, but not with Spa33^C^, which is the same result that was observed for MxiK (**Suppl. Fig. S9**). This is an interesting observation in that it suggests that two proteins found at distant sites within the SP are able to interact with the same component, Spa33^FL^, within the Spa33 heterotrimer. Nevertheless, this does not rule out the possibility that one or both proteins interact with the two SPOA domains of the Spa33 heterotrimer since Spa33^FL^ contains both a SPOA1 and a SPOA2 domain, much like what was described for *Salmonella* SctQ (Notti et al., 2015; McDowell et al., 2016). In line with this, Notti et al. were able to solve the structure of a complex consisting of the two SPOA domains from *Salmonella* SctQ with a fragment of SctL (MxiN homologue) (Notti et al., 2015). Thus, it is interesting that our two-hybrid results did not reveal an interaction with the Spa33^C^ dimer. Perhaps this is because this dimer is generated from two copies of SPOA2 rather than a hybrid SPOA1/SPOA2 structure.

Unlike MxiK, MxiN can be purified as a recombinant protein and when its CD spectrum is obtained, it is clearly rich in α-helical structure with a thermal unfolding transition at 36 °C, which agrees with the value reported by others (**Fig. 5A** and **Table 3**) (Case and Dickenson, 2018). When purified MxiN and Spa33 were mixed (with MxiN being in excess) and the resulting complexes separated by size-exclusion chromatography, MxiN was found to purify in two populations. In addition to MxiN alone (which eluted as the smaller of two eluting peaks of protein by SEC), MxiN also co-purified with the Spa33 complex (**Fig. 3** and **Suppl. Fig. S11**). As with MixK-Spa33, the CD spectrum of the Spa33-MxiN complex was a combination of α-helical structure (contributed by MxiN) and β-structure (contributed by Spa33). When the CD signal at 222 nm was monitored as a function of temperature, the Tm of the major transition was found to be 39 °C (**Fig. 5B** and **Table 3**), indicating that Spa33 has at least a minor stabilizing effect on MxiN. However, a second transition at 62 °C suggests that a minor population of the MxiK-Spa33 complex, this complex may dissociate at the lower transition temperature with the freed MxiN then unfolding with what appears to be a slightly higher Tm than the MxiN alone. The second transition fairly accurately reflects the thermal unfolding reported above for the stable Spa33 heterotrimer (∼60 °C). This prompted us to assess the strength of the MxiN-Spa33 interaction using biolayer interferometry (BLI). MxiN possessing an N-terminal His_6_-tag was immobilized to BLI sensors via bound Ni^2+^ to allow for monitoring the association and dissociation of the wild-type Spa33 heterotrimer using BLI (**Fig. 6** and **Suppl. Fig. S12**). From these data, an equilibrium dissociation constant (K_D_) of 0.55 µM was obtained, suggesting a specific and moderately strong interaction was occurring between them.

**Figure 5.**
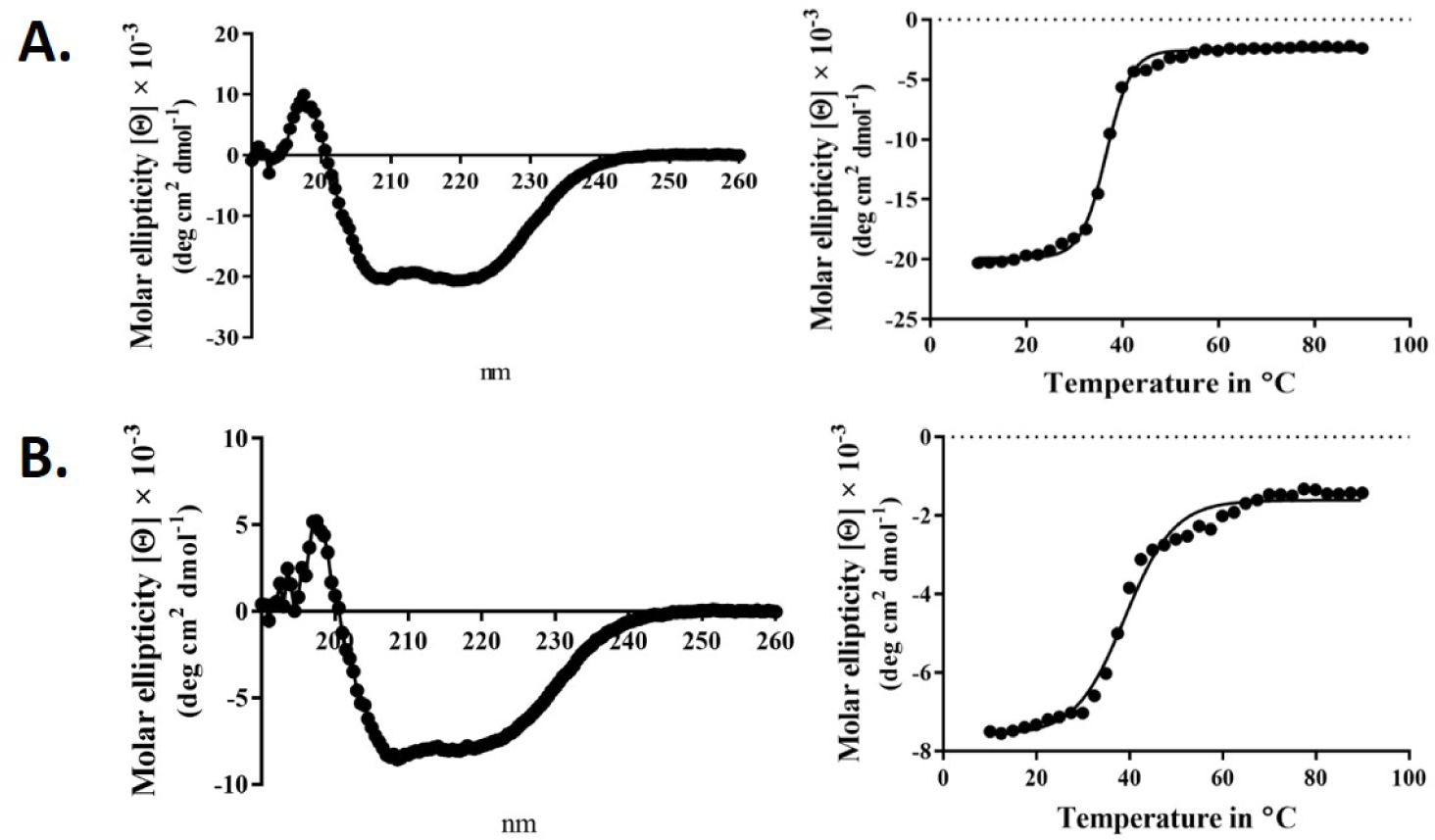
CD spectra from MxiN and the Spa33-MxiN complex. The secondary structures and thermal stabilities of MxiN and its complex with the wild-type Spa33 heterotrimer were analyzed by CD spectroscopy. **A**) The CD spectrum of MxiN alone shows two minima that indicate a rich α-helical structure (**left**). The thermal stability for MxiN was determined by plotting the molar ellipticity at 222 nm as a function of temperature. A single transition was observed giving a Tm value of 36.63 °C (**right**). **B**) The CD spectrum for the MxiN-Spa33 complex also had two minima with the stronger minimum at 208 nm, indicating a greater β-structure content than for MxiN alone (**left**). The thermal stability of the complex was tested by monitoring the molar ellipticity at 222 with increasing temperature. The complex had a major transition with a Tm value of 38.99 °C, however, a minor second transition was seen at about 60 °C (**right**). R^2^ values for these unfolding curves was >0.985.

**Figure 6.**
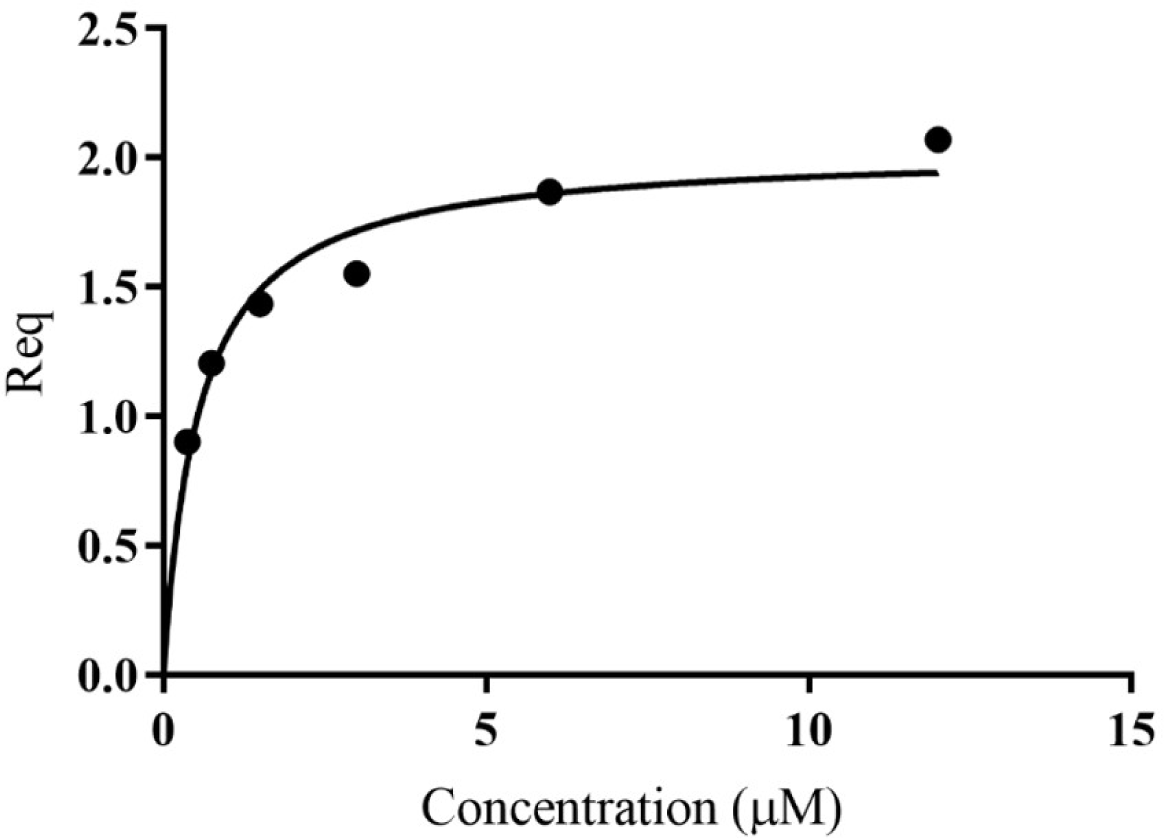
Quantification of the *in vitro* binding of Spa33 to MxiN using biolayer interferometry (BLI). MxiN with an N-terminal His_6_-tag was immobilized to sensor tips to measure its ability to bind to wild-type Spa33 heterotrimers. The tips were immersed in different concentrations of Spa33 (lacking a His_6_-tag) to monitor association and dissociation rates (see **Supplemental Materials**). The K_D_ was determined based on the equilibrium reaction (Req) at different Spa33 concentrations. The K_D_ value was determined to be 0.55±0.091 µM, which is in agreement with the measured association and dissociation rates. The plotted data were generated using GraphPad prism.

**Figure 7.**
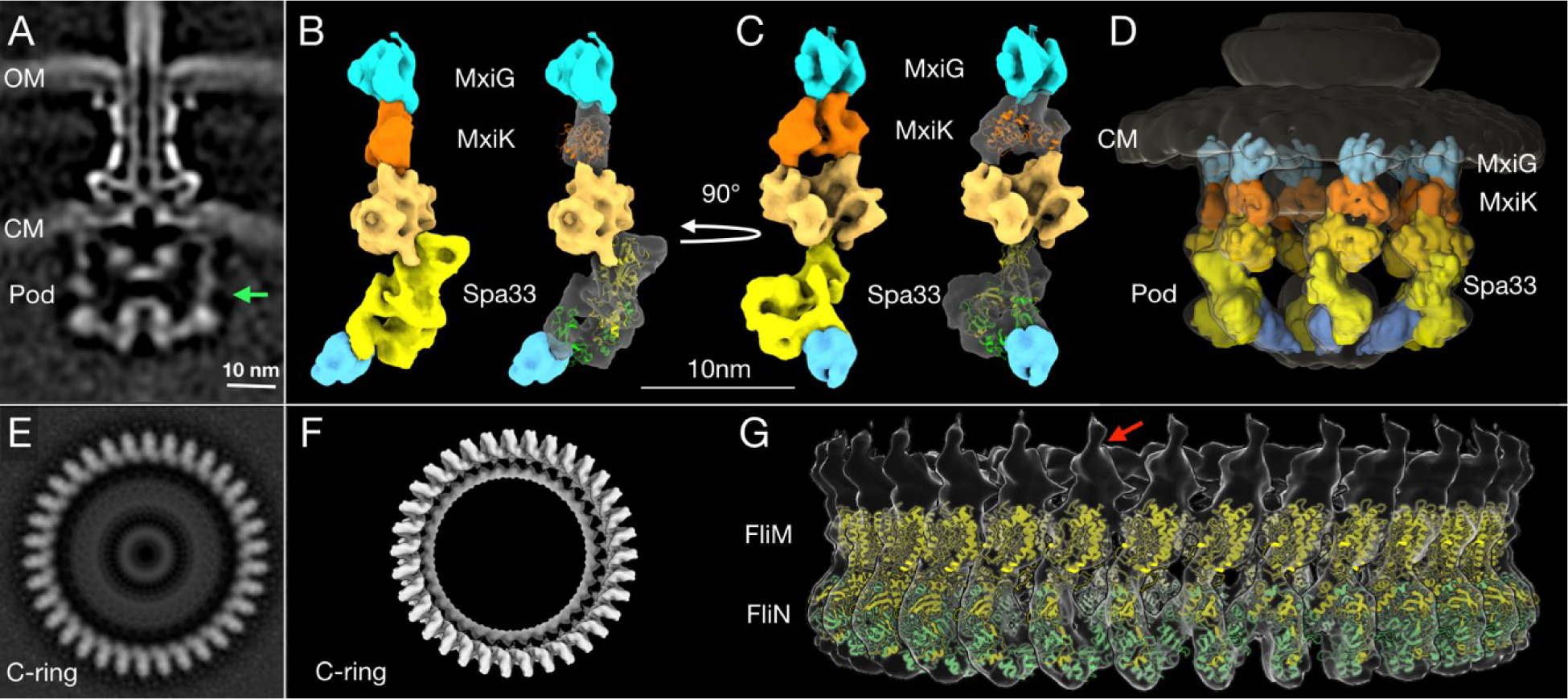
High-resolution structure and proposed model of the *in situ* SP pods from *Shigella*. A) A cross-section image of the subtomogram average structure of the injectisome shows (from **top** to **bottom**) the needle, basal body and SP. The two bacterial membranes are indicated by OM (the outer membrane) and CM (the cytoplasmic membrane). The green arrow indicates the right side of the pod region that was used for the additional refinement process to improve its resolution. **B)** A 3D surface rendering image of the refined pod structure is shown. From the top down, the refined pod structure associates with the cytoplasmic domain of MxiG (blue) and consists of MxiK (orange) and multiple copies of Spa33 (different shades of yellow), which also associate with MxiN (cyan at bottom) (Hu et al., 2015; Tachiyama et al., 2019). The crystal structure of MxiK’s homologue from *P. aeruginosa*, SctK (Muthuramalingam et al., 2020) was fit into the density beneath MxiG (**right**). Two copies of the Spa33 heterotrimer model were fit into the bottom part of Spa33 region in the refined pod structural image (**right**). Because this is a preliminary model, some regions did not fit precisely into the cryo-ET image (see **Suppl. Fig. S15**). The top of the proposed Spa33 density forms a globular structure that interacts with MxiK. **C)** The same images are shown rotated 90゜ relative to **Panel B**. This angle shows the single pod with a view that would be on the inside of the SP cage. **D)** The refined pod structure was fit back into a 3D surface rendering of the complete SP. This image shows the six pods forming the cage-like structure of the SP. **E)** A bottom view of a subtomogram average structure from the *Vibrio alginolyticus* flagella C-ring (Carroll et al., 2020). The C-ring is composed FliM and FliN, which are homologous to Spa33 in the pod assembly. **F)** A surface rendering of the C-ring with its 34-fold symmetry (EMDB-21837) was generated using the UCSF Chimera (Pettersen et al., 2004) and ChimeraX (Goddard et al., 2018) software. **G)** A side view of the 3D surface rendering of the C-ring with a model of a full-length FliM (yellow) and three copies of FliN (green) from I-TASSER (Roy et al., 2010; Yang et al., 2015; Carroll et al., 2020) is shown. The red arrow indicates a single unit of the C-ring structure for comparison with the refined single pod structure in **Panels B** and **C**.

### 3.5. Examination of the Spa33-containing complexes by dynamic light scattering (DLS)

From the CD data on the Spa33-containing complexes and individual proteins presented above, a great deal of information can be obtained regarding secondary structure content and thermal stability. To begin putting these data into context regarding the solution properties for these proteins and complexes, we carried out dynamic light scattering (DLS) studies. DLS allows us to determine the relative sizes of the proteins and their complexes, along with ascertaining the influence that complex formation has on their relative sizes. It also lets us determine whether the complexes are of discrete sizes or are able to nonspecifically form larger species. Furthermore, we obtained the zeta-potential in each case to determine the effect of complex formation on the electrical potential of the particles being examined. As expected, Spa33^C^ is smaller than the wild-type Spa33 heterotrimer (**Table 3**) with hydrodynamic diameters (Dh) of 6 and 13 nm, respectively. The size of the Spa33 complex is more than twice the size of the Spa33^C^ dimer, which is consistent with the compared masses of the two. Spa33^FL^ has been reported to be 35.6 kDa and each Spa33^C^ was reported to be 11.6 kDa based on mass spectrometry analysis (McDowell et al., 2016). This means the heterotrimer would have a mass close to 58.8 kDa relative to the Spa33^C^ dimer’s mass of 23.2 kDa. Nevertheless, the Dh of the heterotrimer is smaller than what might be expected for the entire pod structure based on previous reports (Hu et al., 2015). This is consistent with there being multiple copies of Spa33 or the Spa33-containing complexes within each pod as has been suggested by others (Diepold et al., 2015; McDowell et al., 2016).

Interestingly, the Spa33-MxiK complex has a Dh of 10 nm, which is smaller than that of the Spa33 heterotrimer (**Table 3**), however, the purified Spa33-MxiK complex has all three of the expected components (Spa33^FL^, Spa33^C^ and MxiK) present (**Fig. 3**). Furthermore, the Spa33-MxiK complex has a dominant transition for thermal unfolding (**Fig. 4**) and this Tm is distinct from that shown for the Spa33 heterotrimer. This suggests that much of the Spa33-MxiK complex acts as a single entity. The absence of multiple complex forms is also evident from the DLS scans when either particle numbers (**Suppl. Fig. S13**) or intensity (not shown) are examined. This also suggests that the Spa33 component undergoes a significant conformational change when in a complex with MxiK and that the overall particle has a more compact globular structure than the elongated Spa33 heterotrimer. The Dh for MxiK could not be determined because this protein is not soluble on its own in solution and could not be purified. Nevertheless, initial reports of the interface between MxiK and the Spa33 pods using cryo-ET imaging suggest an intimate interaction between the two (Tachiyama et al., 2019).

In contrast to MxiK, MxiN can be purified and it was found to have a Dh of almost 10 nm, and this is consistent with its ability to form dimers that become the elongated spokes connecting the pods with Spa47 (Hu et al., 2015). When incubated with Spa33 *in vitro*, the resulting complex has a Dh of 18.5 nm and this is what might be expected following the association of the two structures (**Table 3**). This behavior is distinct from that of the Spa33-MxiK complex, but like the Spa33-MxiK complex there is single-sized species that is formed (**Suppl. Fig. S13**). From the two-hybrid, CD and DLS data, we propose that Spa33 forms two distinct complexes that are stable in solution, one incorporating MxiK and one incorporating MxiN. In each of these complexes, it appears that the interaction with Spa33 occurs through the Spa33^FL^ moiety. Whether this means that the interactions occur through the N-terminal portion of Spa33 is not yet clear, however, Spa33^C^ alone has not been found to be directly responsible for the protein-protein interactions giving rise to these complexes though it does contribute to their stabilization.

In addition to Dh, we monitored the surface charge properties (zeta-potential) of these proteins and their complexes. Spa33 and Spa33^C^ both have a negative surface electric potential (Spa33 = −8.87 nV) while MxiN has near neutral surface charge. The fact that the Spa33-MxiN complex has a zeta-potential of −5 mV suggests that this interaction masks some of the surface electric potential of Spa33. In contrast, the complex with MxiK only reduces the negative zeta-potential of Spa33 by about 1.5 mV. While we do not know the zeta-potential of MxiK alone, this suggests that there are significant differences in how it interacts with Spa33 in comparison to MxiN.

### 3.6. Visualization of the sorting platform pods by cryo-electron tomography (cryo-ET) imaging

Bzymek et al. first described the independent expression of a SctQ C-terminal domain in *Yersinia* (Bzymek et al., 2012). Notti et al. then identified SPOA domains and described their relationship to flagellar FliN in their work in the *Salmonella* system (Notti et al., 2015). For the *Shigella* system, the two SP structures that have been solved thus far include the SPOA domain of Spa33, which forms a dimer (the Spa33^C^ homodimer) (McDowell et al., 2016) and the Spa47 ATPase (Burgess et al., 2016). While MxiK and MxiN have not had their high-resolution structures solved. Our group recently reported the crystal structure of the MxiK homologue from *P. aeruginosa*, SctK (Muthuramalingam et al., 2020). The latter has a unique fold and is the first structure solved for any protein in the SctK family. No structures are available for the virulence-related T3SS Spa33/SctQ heterotrimer or any MxiN family member. There are, however, structures available for FliN (**Suppl. Fig. S3**) and FliM (**Suppl. Fig. S14**) from *T. maritima* (Brown et al., 2005). We chose to use these as surrogates for modeling the Spa33 heterotrimer into the pod densities of the SP. FliM-FliN is a heterotetramer (Paul et al., 2006) that has been proposed to be the structural equivalents of the Spa33 heterotrimer. In such a model FliM would be equivalent to the N-terminal portion of Spa33, despite there likely being significant structural differences between them, and the FliN trimer would represent the three copies of Spa33^C^ within the overall Spa33 complex (comprising two copies of Spa33^C^ with a third copy of the SPOA domain contributed by one of the two tandem SPOA domains at the C-terminus of Spa33^FL^). Using this and the flagellar C-ring of *Vibrio alginolyticus* (Carroll et al., 2020) as a guide for generating a model of the Spa33 heterotrimer and how such Spa33 complexes might form an interface, we could insert two copies of the modeled Spa33 homotrimer into a newly generated high-resolution image of a single SP pod from the *Shigella* injectisome (**Fig. 7**). The fit agrees with the data on the size and shape of wild-type Spa33 when it is alone and/or incorporated into complexes with MxiN and MxiK (**Table 3** and **Fig. 7C-D**).

The orientation of the Spa33 units in our model is different from what is seen in the flagellar C-ring of *V. alginolyticus*. Within the flagellar C-ring, multiple copies of the FliM-FliN complex align side by side to generate a contiguous ring (**Fig. 7E-G**), however, the Spa33 heterotrimer would be oriented differently so that it is nearly on its side to form a complex with another Spa33 complex that lies above it (**Fig. 7B-C**). This would then give rise to six distinct pod entities evenly placed around the central ATPase to which it is connected via MxiN. Meanwhile, the upper Spa33 heterotrimer is shaped slightly differently in that it forms a tight complex with MxiK whose density appears more globular than of MxiN in the *in situ* SP (**Fig. 7D**). The Spa33 component of the Spa33-MxiK complex in this case undergoes a conformational change as it interfaces with MxiK and this gives rise to a more compact globular structure compared with the Spa33-MxiN complex. This fits with the observation that the Dh of the Spa33-MxiK, which is slightly smaller than that of the Spa33 heterotrimer alone. The structure of SctK from *P. aeruginosa* is fit into the density occupied by MixK where it forms an intimate interface with the upper Spa33 heterotrimer (**Fig. 7C-D** and **Suppl. Fig. S15**). Thus, the Spa33 moiety within the Spa33-MxiN and Spa33-MxiK complexes are proposed to undergo conformational changes that give rise to two distinct complexes within the SP – one for interacting with the IR via MxiG^C^ and one for interacting with the Spa47 ATPase via MxiN.

## 4 Discussion

This work provides a detailed biochemical and biophysical exploration of the *Shigella* T3SS sorting platform (SP). The component protein interactions described here have been proposed before based on different types of analyses, including two-hybrid analysis and pull-down assays (Jouihri et al., 2003; Morita-Ishihara et al., 2006; Johnson and Blocker, 2008), however, the data presented here and related cryo-ET studies are making significant contributions to our detailed understanding of the structures and interfaces that are important for SP assembly (Hu et al., 2015; Hu et al., 2017).. The pods are the major component of the *Shigella* injectisome SP. A difficulty encountered in focusing on these structures *in situ* is their apparent flexibility and dynamics within the active SP. Nevertheless, based on our findings, we can propose a model for the assembly of the Spa33-containing pods and speculate as to how these individual sub-assemblies contribute to active type III secretion in *Shigella*. It is accepted that dynamic processes within the SP are critical for recruitment of secretion substrates to the injectisome and that the SctN ATPase has an essential role in presenting these substrates to the export gate (Diepold et al., 2017). As part of this process, SctN has been proposed to act as a rotary motor (Majewski et al., 2019) in guiding substrates to the export gate, perhaps via the central stalk formed by SctO (Spa13 in *Shigella*). Dielpold *et al*. reported that *Yersinia* SctQ, a Spa33 homologue, is present in more than 20 copies within the injectisome SP and that it is in dynamic exchange with a cytoplasmic population during active type III secretion (Diepold et al., 2017). The details of how this dynamic exchange occurs remains to be determined and how this applies precisely to the *Shigella* system is far from clear.

It was recently reported by Bernal *et al*. that soluble complexes consisting of the *Salmonella* SP components SctQ^FL^, SctQ^C^, SctL and SctN could be assembled and the organization of this complex could be determined using small-angle X-ray scattering (SAXS) (Bernal et al., 2019). As with the Spa33 complex, SctQ^FL^ was made soluble by the presence of SctQ^C^. The SctQ heterotrimer could associate with a SctL dimer in the same way that Spa33 and MxiN interact. They further showed that the SctQ-SctL complex could associate with the SctN ATPase. This is similar to the work of Notti et al. (Notti et al., 2015) who could purify a complex consisting of SctQ, SctL and SctN. We did not attempt to generate a Spa33-MxiN-Spa47 complex, however, this is a relationship that will need to be explored as this work continues. We also found that in addition to MxiN, the Spa33 heterotrimer could associate with MxiK, however, this required that the wild-type *spa33* and *mxiK* be co-expressed prior to purifying the resulting complex. This association has a stabilizing (and solubilizing) effect on MixK and, based on the poor solubility of MixK, it seems likely that the MxiK that exists in the *Shigella* cytoplasm is in fact part of a Spa33-MxiK complex that for the most part behaves as a single folded unit that is highly stable. Moreover, the Spa33-MxiK complex has biophysical properties (thermal stability and hydrodynamic radius) that are distinct from those of the Spa33-MxiN complex, which can be assembled *in vitro* from purified components.

We have demonstrated that MxiK is an essential SP component that acts as an adaptor between the SP and the cytoplasmic domain of the IR protein MxiG. In the absence of MxiK, the SP is completely absent from the injectisome. Meanwhile, MxiN forms radial spokes that emanate from a central hub that is the Spa47 hexamer (Hu et al., 2015). In the absence of MxiN, remnants of the pods are still visible and these are the result of the interactions between MxiG, MxiK and Spa33 (**Suppl. Fig. S2**) (Hu et al., 2015). This unique observation indicates that a sub-assembly consisting of MxiK and Spa33 can be incorporated into an otherwise absent SP. That cannot be said of MxiN-Spa33 subassemblies and this is why we proposed that the Spa33-MxiK and Spa33-MxiN complexes can form and behave independently in the *Shigella* cytoplasm. Furthermore, it appears that the Spa33-MxiK units can be associate with the injectisome basal body in the absence of MxiN (**Suppl. Fig. S2**).

From the data presented here and elsewhere, we propose that the Spa33-MxiK and Spa33-MxiN assemblies are physiologically relevant structures with important roles in the mechanisms governing type III secretion in *Shigella*. These data are also consistent with the large pod structures of the SP being formed from two copies of the Spa33 heterotrimer with an architecture that is altered from the interactions that give rise to a contiguous flagellar C-ring. Thus, in view of the dynamics involving SctQ exchange between the SP and cytoplasm, we propose a model in which Spa33 heterotrimers act at two different locations during the recruitment of secretion cargo. Spa33-MxiK complexes are thus proposed to be required for communication between the injectisome IR and the SP while Spa33-MxiN complexes are active in ushering secretion substrates to the SP where they can then be acted upon by Spa47 for chaperone removal and presentation to the export gate (Akeda and Galan, 2005). It is also possible that the Spa33-MxiN complex is interacting with Spa47 prior to incorporation (with secretion substrate) into the SP based on independent work by Notti et al. and Bernal et al. (Notti et al., 2015; Bernal et al., 2019). The export gate then harnesses the proton-motive forces to actually fuel the secretion process (Shen and Blocker, 2016). It is also likely that the Spa33-MxiK complex is in exchange with a cytoplasmic pool of similar complexes. Interestingly, this complex has the potential to act independent of the Spa33-MxiN complex in interacting with the IR (Hu et al., 2015), perhaps to act in guiding Spa33-MxiN-effector (and possibly Spa47) complexes to the SP where there are up to five other pod complexes comprised of MxiK, Spa33 and MxiN in association with the Spa47 hub. Such a model fits with the cryo-ET based structural fits depicted in **Figure 7**, which are modeled based on the similarities between the now solved structure of *Pseudomonas* SctK and a model of the Spa33 heterotrimer based on similarities with the flagellar C-ring components FliM and FliN (**Suppl. Fig. S15**). While perhaps speculative, this model fits with the data presented. A great deal of work remains to fully understand the events occurring within the SP, however, complementary molecular, biophysical and imaging techniques are beginning to shed light on how the full injectisome functions. Perhaps further revelations will become known when we are able to manipulate the *Shigella* T3SS so that it adopts specific secretion states.

## Conflict of Interest

Author Ryan Skaar was employed by the company NanoBio Designs, LLC during the writing of this manuscript. The author Michael L. Barta was employed by the company Catalent Pharma Solutions during the writing of this manuscript. The remaining authors declare that the research was conducted in the absence of any commercial or financial relationships that could be construed as a potential conflict of interest.

## Author Contributions

W.D.P. was responsible for conceptualization, funding acquisition, project administration, writing portions of the original draft and supervision of research staff; S.T. was responsible for conceptualization, investigation, methodology and writing portions of the original draft; R.S. was responsible for two-hybrid analyses, portions of the biophysical analyses and critical reading of the manuscript; Y.C. was responsible for the cryo-ET analyses and refinements; B.C. was responsible for data analysis and modeling; M.M. was responsible for data curation, formal analysis, investigation, methodology and critical review of the manuscript; S.K.W. was responsible for conceptualization, formal analysis, investigation, methodology, visualization and writing portions of the original draft; M.L.B. was responsible for generating fusion protein mutants and portions of the contact-hemolysis experiments; W.L.P was responsible for manuscript editing and writing; J.L. was responsible for conceptualization, overseeing the cryo-ET imaging studies and data analysis and interpretation.

## Funding

This research was funded by the National Institute of Allergies and Infectious Disease awards NIAID R01 AI123351 and R21 AI146517 to WDP.

## Contribution to the Field

We show here that the Spa33-containing pods of the *Shigella* T3SS sorting platform are formed by interactions between multiple Spa33 heterotrimers. Furthermore, distinct Spa33-containing complexes having different biophysical properties are formed with the adaptor protein MxiK and the radial spoke protein MxiN. Based on stable complexes and high-resolution cryo-electron tomographic imagining, we propose a dynamic model for the exchange of these two complexes with cytoplasmic pools of the same free complexes that work cooperatively to deliver secretion substrates to the SP for presentation to the injectisome export gate.

---

**Supplemental Figure S1.**
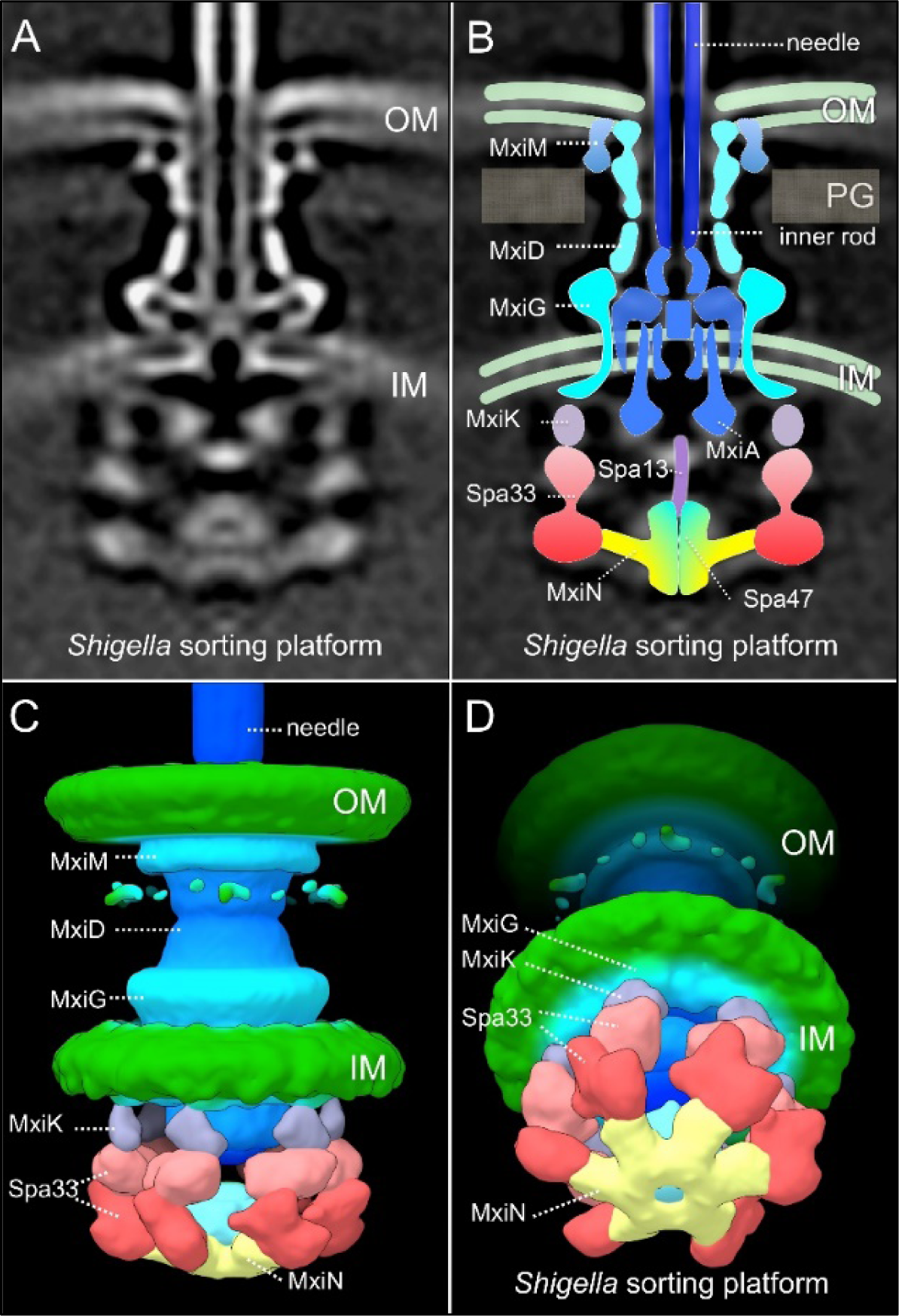
Arrangement of the SP “pods”. A cryo-ET image of the *in situ* wild-type *Shigella* injectisome is shown in **Panel A**. **Panel B** shows the position of the major injectisome components relative to the inner (IM) and outer (OM) membranes and peptidoglycan cell wall (PG) indicated. Based on the cryo-ET image in **A**, a rendering of the injectisome structure is shown in **Panel C** with MxiK (SctK) being the adaptor protein between the Spa33 (SctQ) pods and the cytoplasmic domain of MxiG (SctD) of the IM ring. **Panel D** shows a tilted version of the injectisome that focuses on the SP with the Spa33 densities (red) making up the bulk of the pods. MxiN (SctL) provides the radial spokes that connect Spa33 to the central hub which is the Spa47 (SctN) ATPase. The Spa33 complex consists of what appears to be two large lobes (**Panels B-D**). This figure was originally presented in Hu B, Morado DR, Margolin W, Rohde JR, Arizmendi O, Picking WL, Picking WD, Liu J (2015) Visualization of the type III secretionsorting platform of *Shigella flexneri*. Proc Natl Acad Sci USA 112:1047-1052 (doi: 10.1073/pnas.1411610112).

**Supplemental Figure S2.**
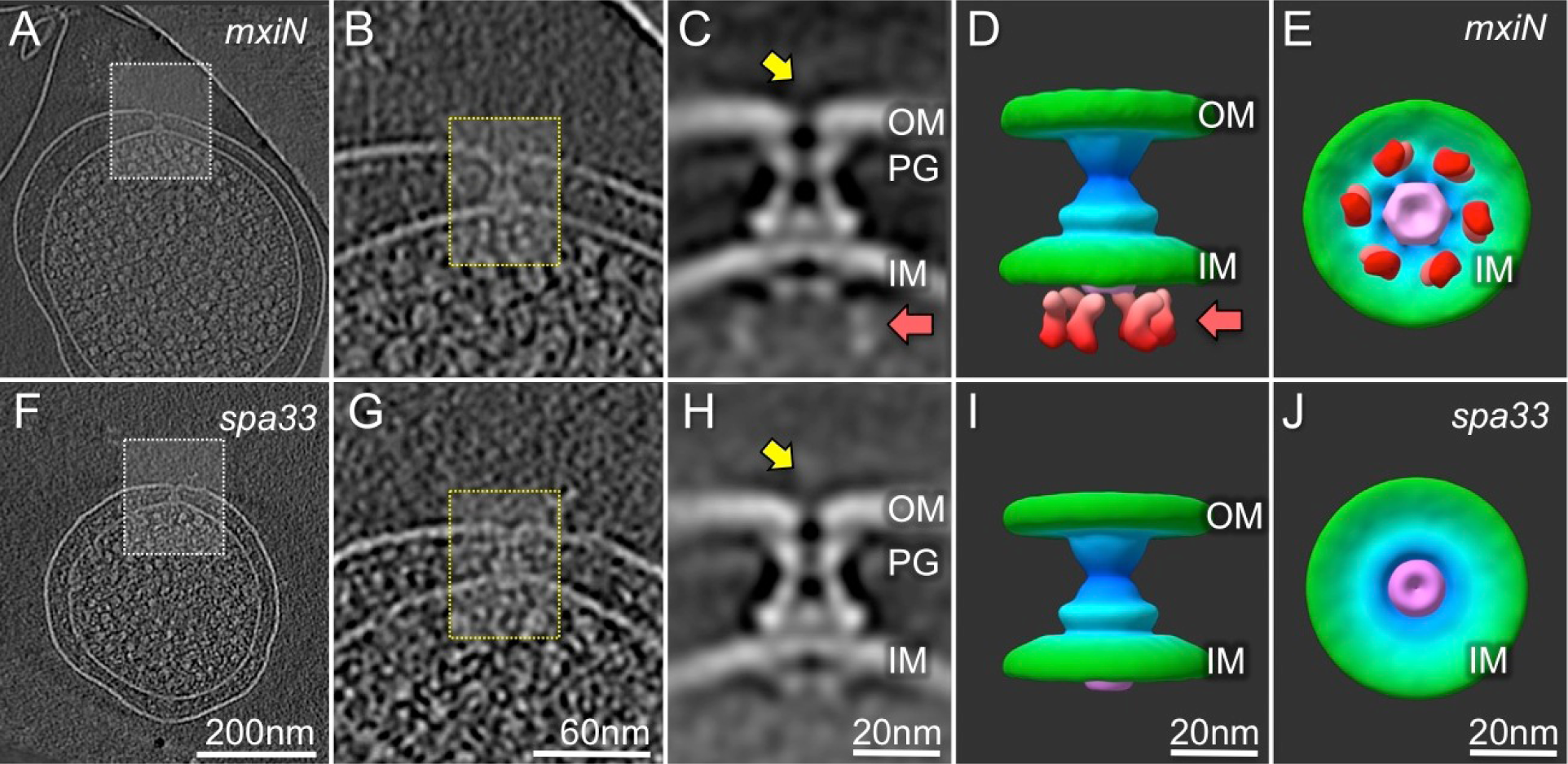
Two injectisome structures from *S. flexneri* minicells lacking MxiN or Spa33. Representative slices of cryo-ET reconstructions of a Δ*mxiN* minicell **(A)** or a Δ*spa33* minicell **(F)**. The corresponding zoomed-in views are shown in **(B)** and **(G)**, respectively. The averaged structure derived from Δ*mxiN* injectisomes is shown in (**C**), together with two 3-D surface views **(D** and **E)**. The averaged structure derived from Δ*spa33* injectisomes is shown in **(H)**, together with two 3-D surface views **(I** and **J)**. This figure was originally presented in Hu B, Morado DR, Margolin W, Rohde JR, Arizmendi O, Picking WL, Picking WD, Liu J (2015) Visualization of the type III secretionsorting platform of *Shigella flexneri*. Proc Natl Acad Sci USA 112:1047-1052 (doi: 10.1073/pnas.1411610112).

**Supplemental Figure S3.**
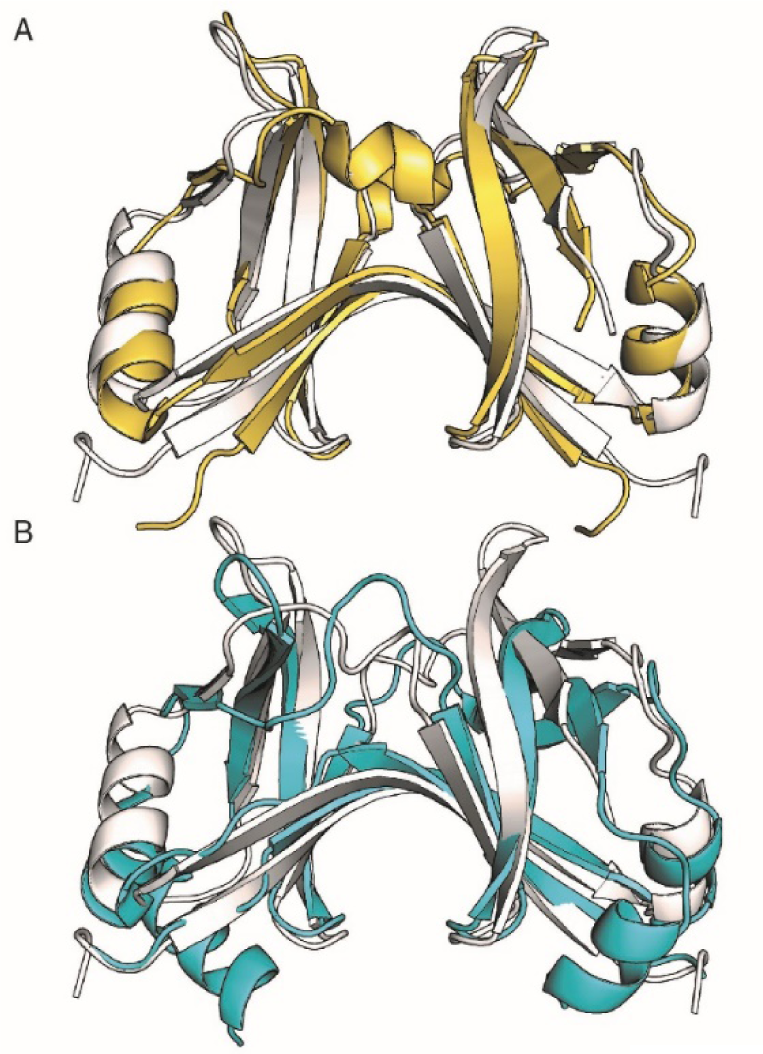
SPOA domain structures. **A)** The crystal structures of the injectisome pod protein components Spa33^C^ (PDB: 4TT9) from *Shigella* (white) and SctQ^C^ (PDB: 4YX1) from *Salmonella* (yellow) are shown (RMSD = 1.4 Å). **B)** The Spa33^C^ structure (white) also compares favorably with the flagellar C-ring protein FliN (PDB: 1YAB) from *Thermotoga maritima* (cyan) (RMSD = 2.1 Å).

**Supplemental Figure S4.**
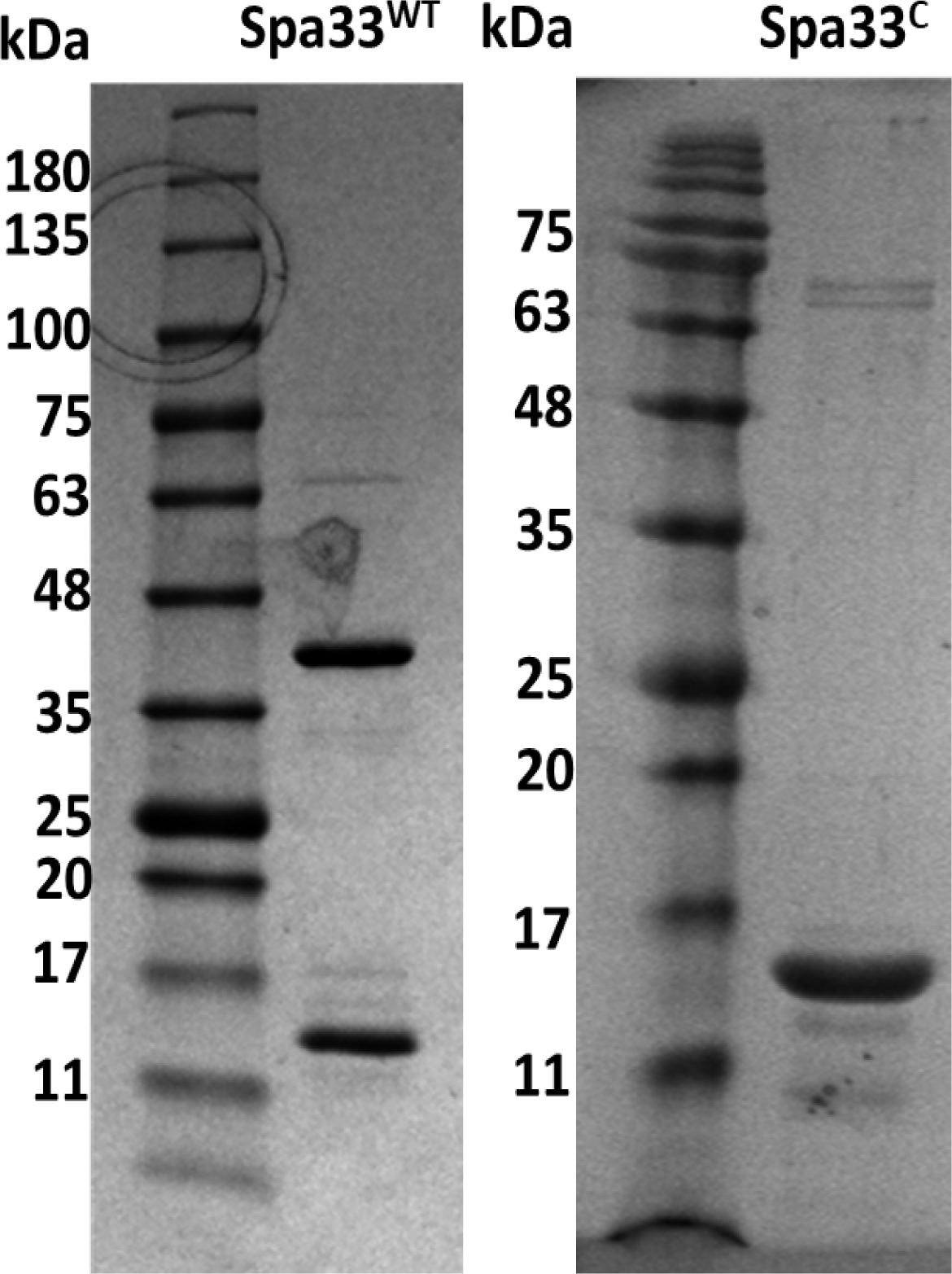
Expression of recombinant Spa33 in *E. coli*. The final purification products were analyzed by SDS-PAGE. A His_6_-tagged version of wild-type Spa33 shows two bands on the SDS-PAGE gel (**left**). The bands at approximately 38 kDa and 13 kDa based on the molecular weight markers are His_6_-tagged Spa33^FL^ and Spa33^C^, respectively. The final purification product of Spa33^C^ is shown in the **right** image with the size being slightly larger than in the **left** due to the presence of a His_6_ affinity tag. Only one dominant band was visible in this case. Molecular weight markers are shown in the first lane of both Coomassie blue-stained gels.

**Supplemental Figure S5.**
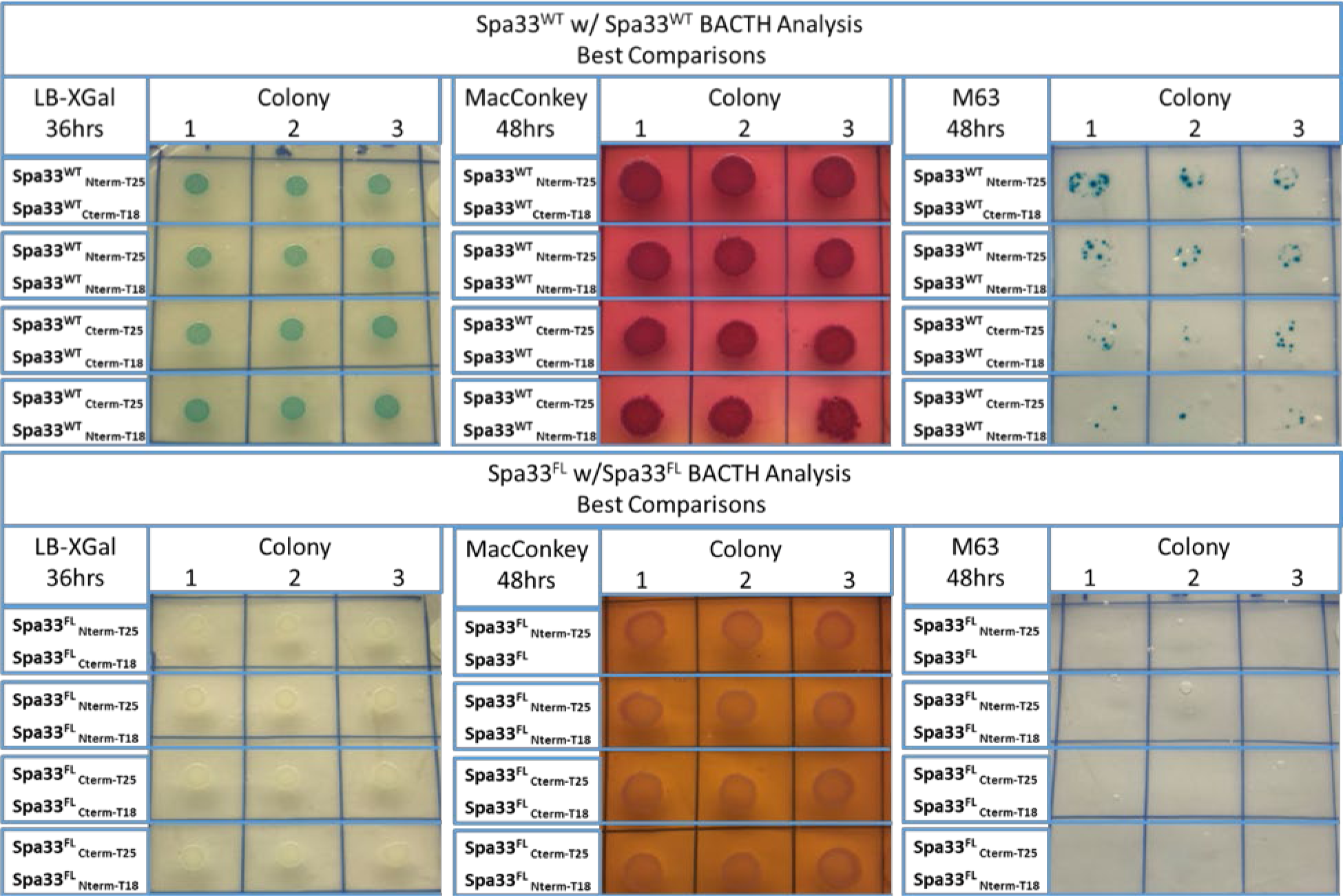
Wild-type Spa33 shows an interaction with wild-type Spa33 while full-length Spa33 (Spa33^FL^) does not show an interaction with another Spa33^FL^ in BACTH analyses. Interaction among components of wild-type Spa33 was tested using a BACTH analysis (**top**). All combination of wild-type *spa33* with the T18 and T25 vectors were co-transformed into *E. coli* BTH101. All bacterial spots had turned blue on X-Gal plates (**left and right**) and red on MacConkey plate (**middle**). While the entire colony did not turn blue on the M63 plates, punctate blue patches were observed within each overall colony. Then the interactions between Spa33^FL^ and a second copy of Spa33^FL^ was tested (**bottom**). In this case, none of the combinations of Spa33^FL^ fused with T18 or T25 gave the color change that would indicate that they interact.

**Supplemental Figure S6.**
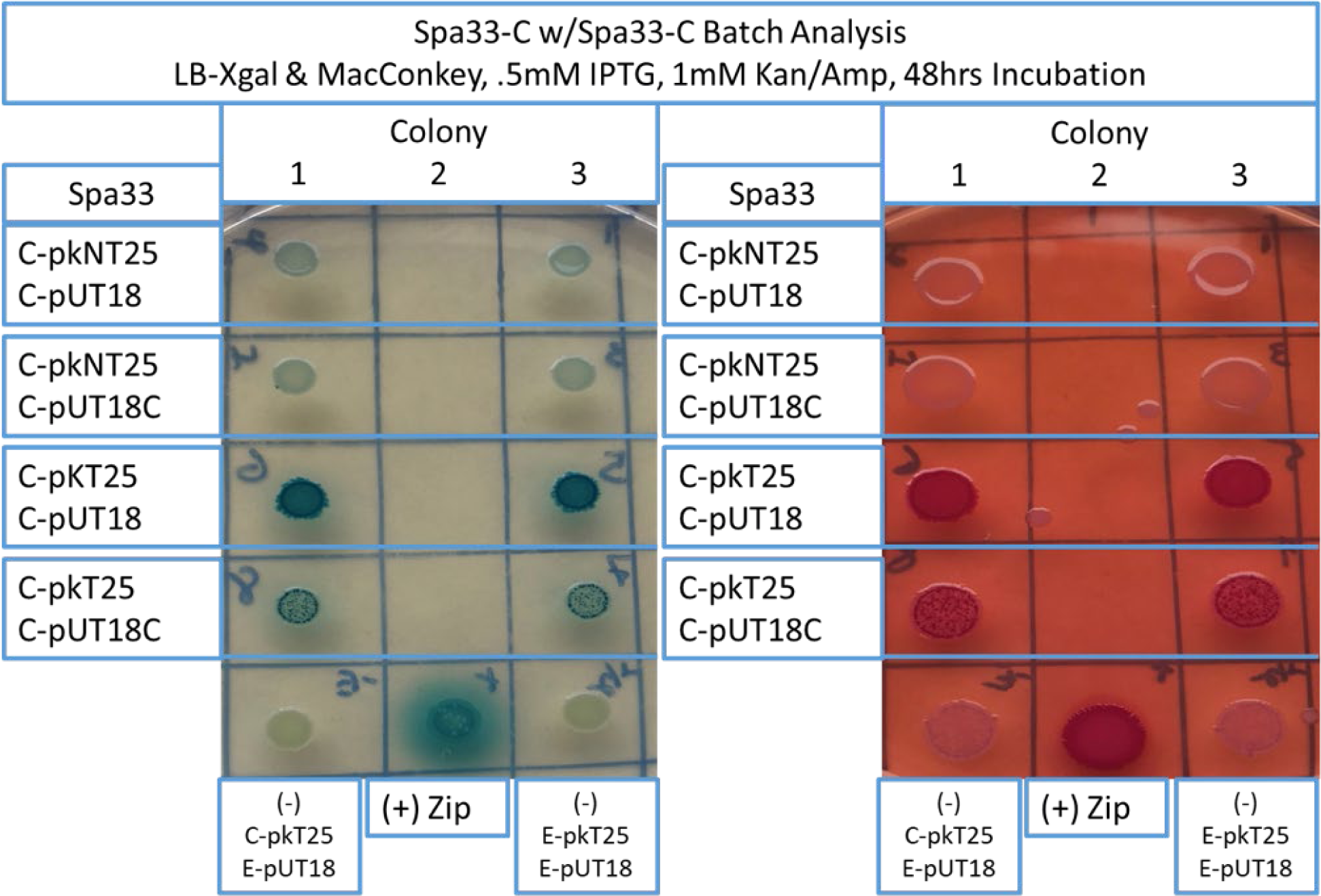
Spa33^C^ interacts with Spa33^C^ in BACTH analyses. Not all combinations of the T18 and T25 vectors with spa33^C^ used to test for interaction for this dimeric protein were positive. However, *spa33^C^* in pKT15 and pUT18 and *spa33^C^* in pKT25 and pUT18C clearly showed color changes on the indicator plates. This experiment confirmed that this two-hybrid analysis can detect the formation of Spa33^C^ dimers.

**Supplemental Figure S7.**
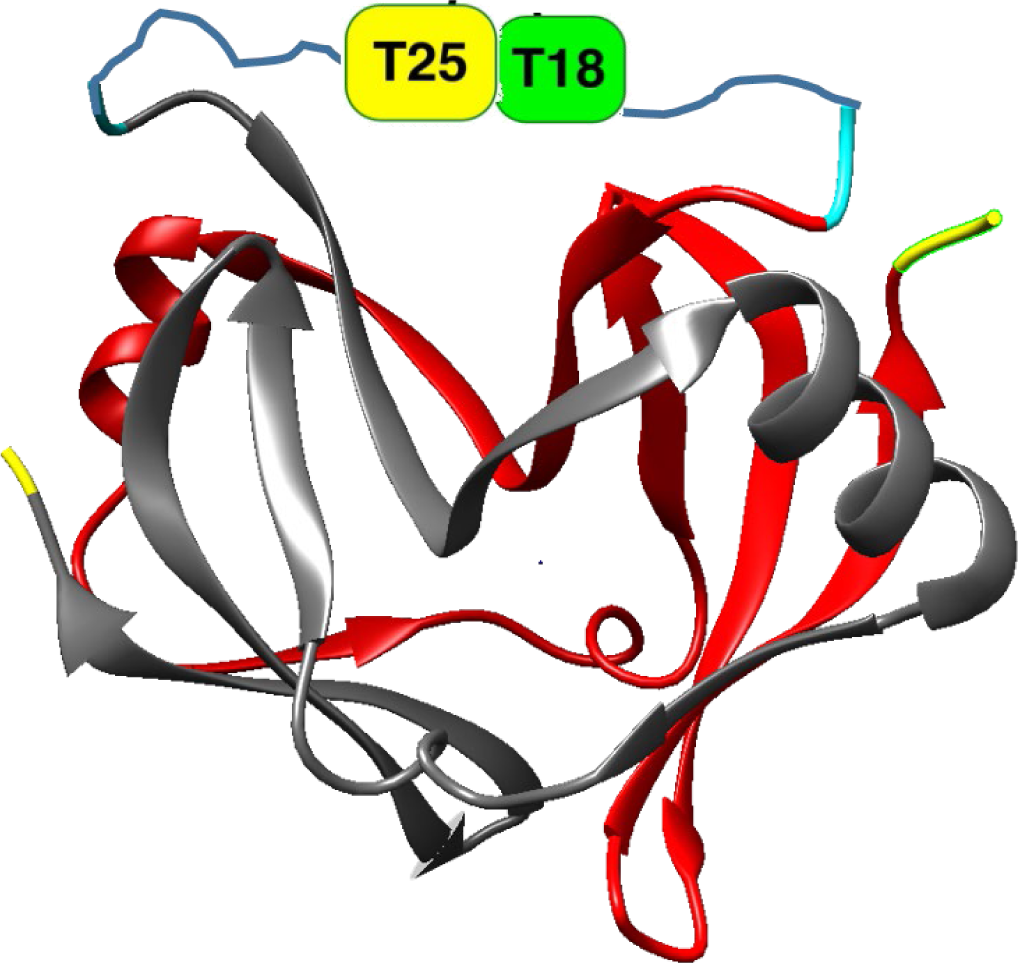
The interacting Spa33^C^ partners must allow for the proper orientation of the CyaA T18 and T25 domains to permit restoration of adenylate cyclase activity. This figure illustrates how two CyaA subdomains, T18 and T25, in Spa33^C^ may be able to interact and restore CyaA activities in *E. coli* BTH101. T18 and T25 subdomains must be localized in the appropriate location and orientation to restore its enzymatic activity during the interaction. This is why not all combinations in Supplemental Figure S5 restored CyaA β-galactosidase activity. (The Spa33 structure is from PDB ID: 4TT9.)

**Supplemental Figure S8.**
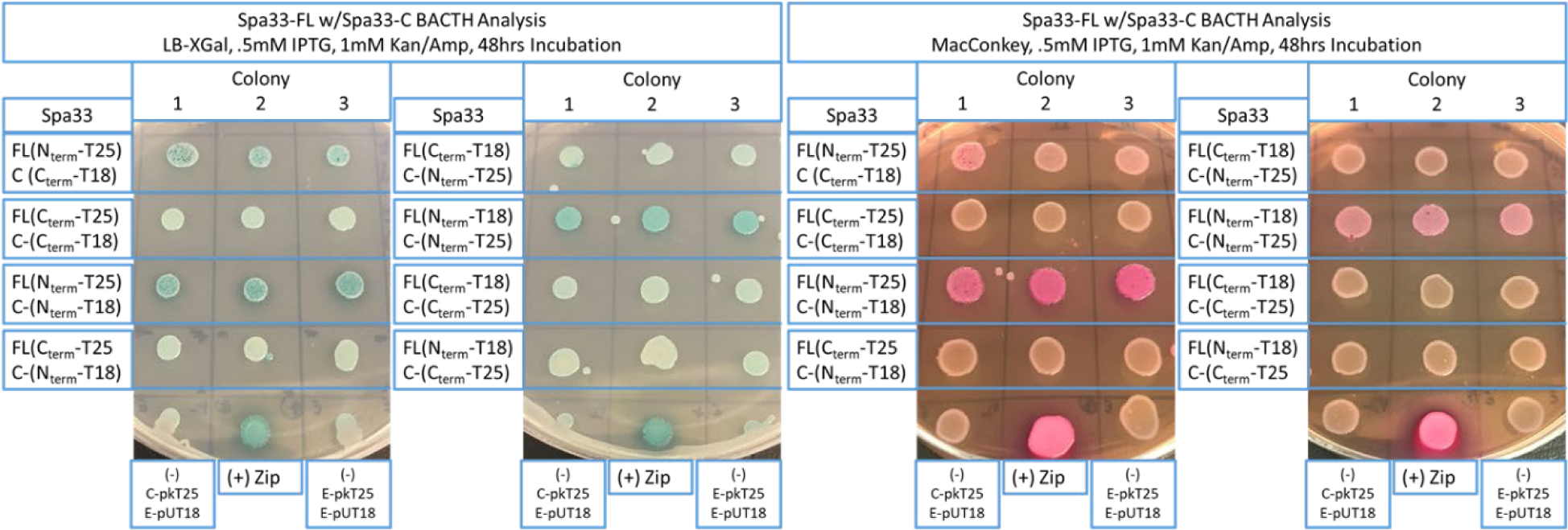
Spa33^FL^ is able to interact with Spa33^C^ in BACTH analyses. All combinations of *spa33^FL^* and *spa33^C^* in T18 and T25 plasmids were co-transformed into *E. coli* BTH101 strain to test their interactions. Some but not all combinations of these fusion plasmids turned blue and red on LB plate with X-Gal and MacConkey plate with maltose, respectively.

**Supplemental Figure S9.**
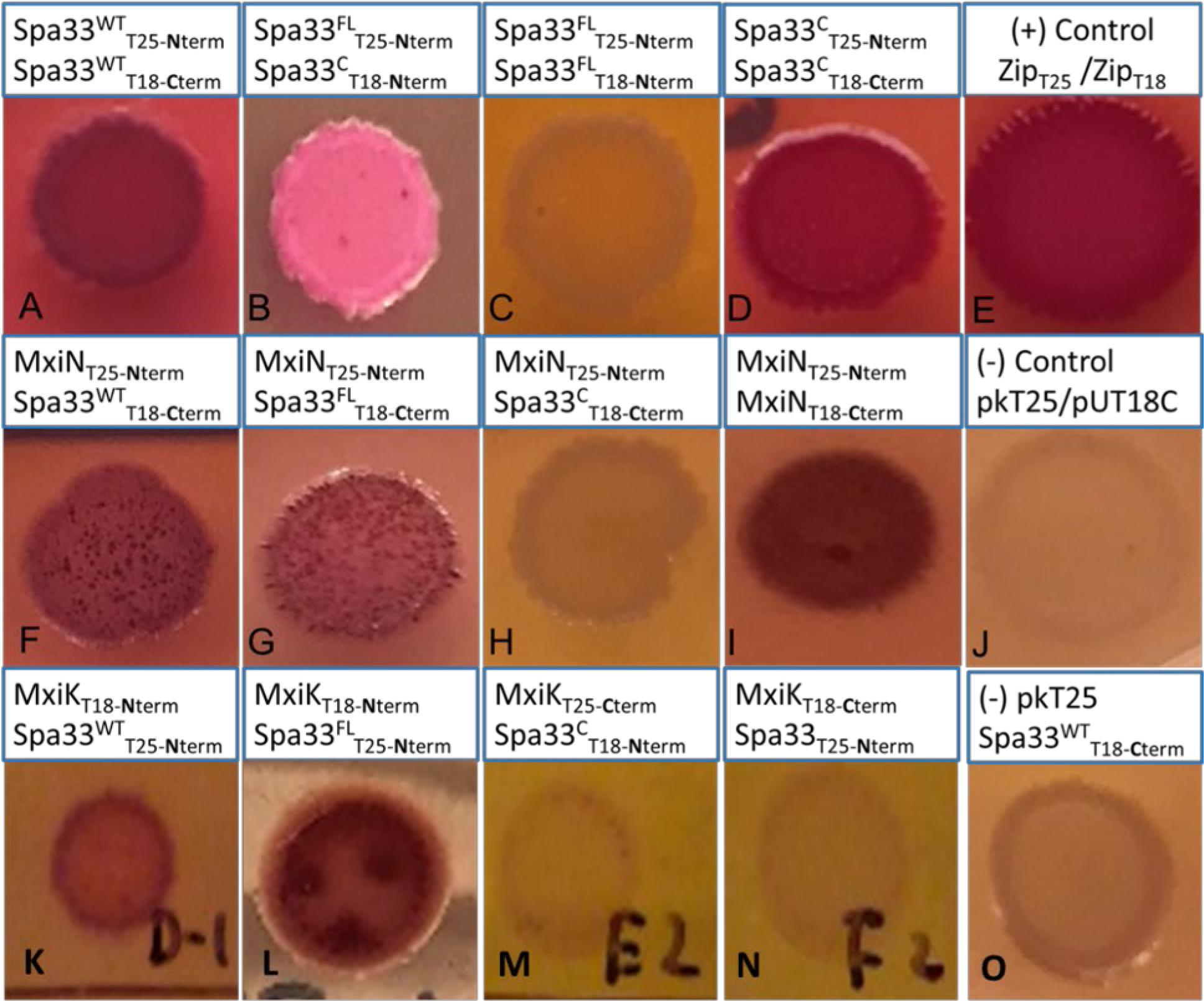
BACTH analysis shows that Spa33 interacts with MxiN and MxiK through Spa33^FL^. As already shown, wild-type Spa33 can interact with itself, Spa33^FL^ can interact with Spa33^C^, Spa33^FL^ cannot interact with itself and Spa33^C^ can interact with itself (**top row**). In contrast, wild-type Spa33 and Spa33^FL^ interact with MxiN, but Spa33^C^ does not interact with MxiN (and MxiN is able to interact with itself) (**middle row**). Likewise, wild-type Spa33 and Spa33^FL^ interact with MxiK, but Spa33^C^ does not (**bottom row**). Not all BACTH plasmid permutations are shown here, however, in no case was any interaction between Spa33^C^ detected in these analyses.

**Supplemental Figure S10.**
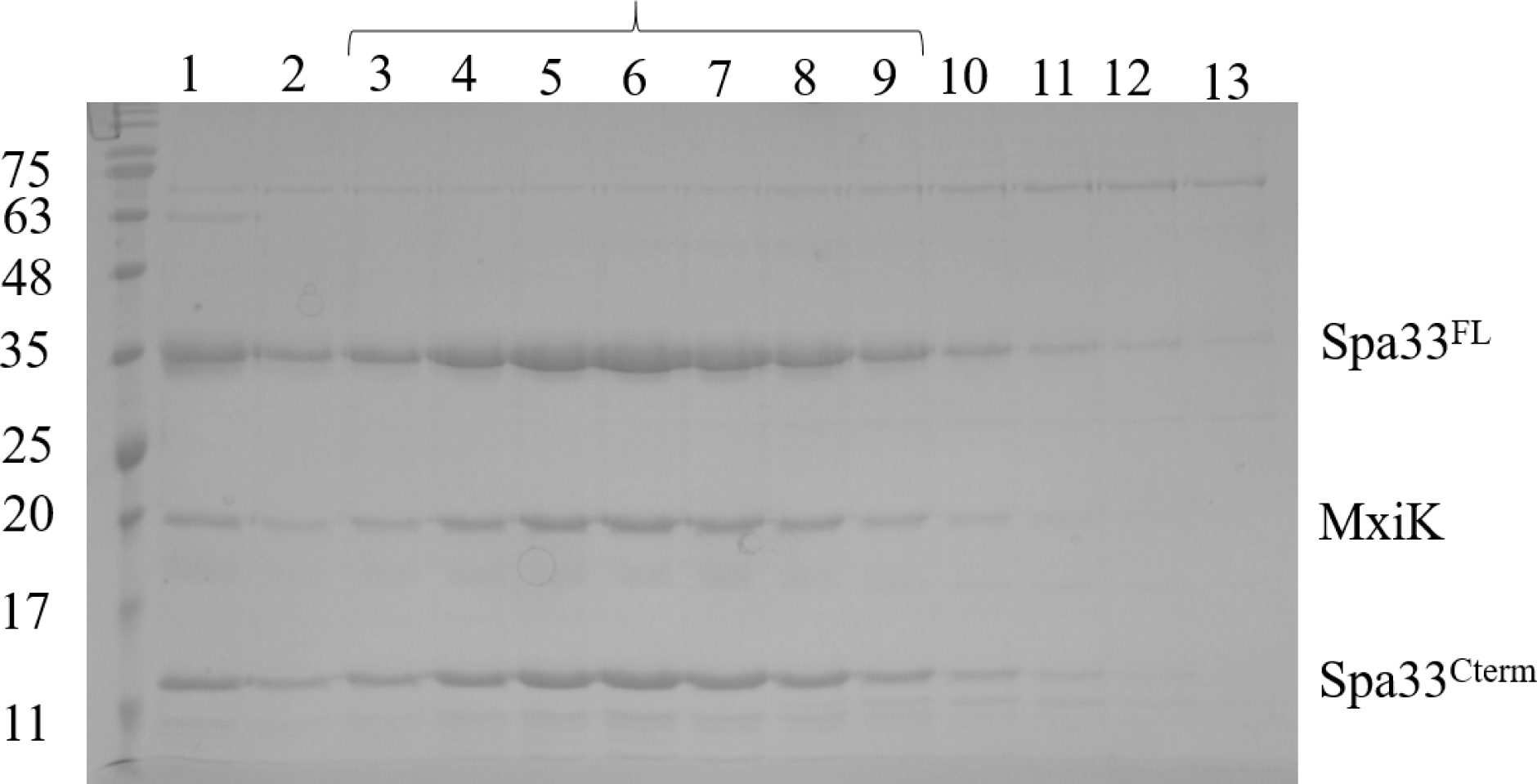
SDS-PAGE analysis of Spa33^WT^-MxiK complex. Wild-type Spa33 and MxiK were co-expressed in *E. coli* and purified on an IMAC affinity column by virtue of a his-tag at the N-terminus of Spa33. The IMAC fractions of the purified complex was then further purified by size-exclusion chromatography (SEC) on a 200pg column with the eluting fractions shown here. Spa33, which is composed of Spa33^FL^ and Spa33^C^, and MxiK were co-eluted from the purification column. The protein components are indicated at the right and fractions 3 to 9 represent the peak of the co-eluting proteins.

**Supplemental Figure S11.**
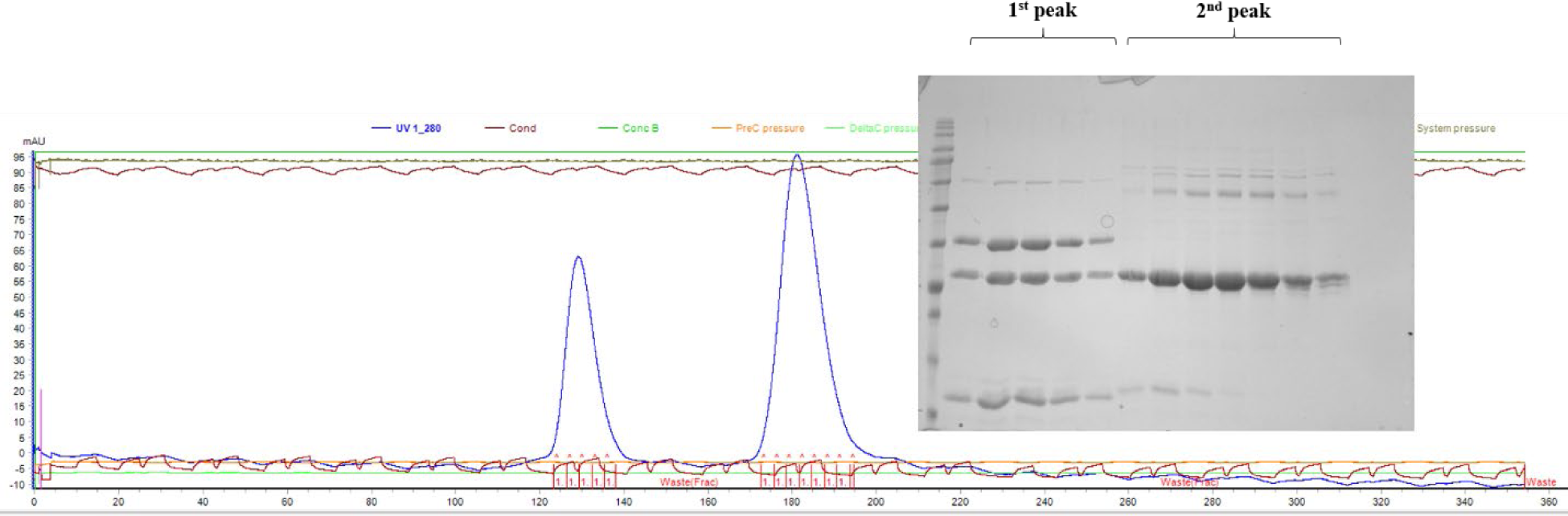
Size-exclusion chromatography profile for the Spa33-MxiN complex (first peak) and residual MxiN (second peak). MxiN and Spa33 were purified by IMAC with their His_6_-tags subsequently removed. The two where then mixed with an excess of MxiN with the protein sample then was then subjected to size-exclusion chromatography. The chromatogram showed two different populations were eluted. SDS-PAGE was then used to analyze the eluted proteins. Fractions from the first (higher molecular weight) peak contained Spa33^FL^, Spa33^C^ and MxiN. The second peak exclusively contained the excess MxiN, indicating all of the Spa33 heterotrimer was associated with MxiN.

**Supplemental Figure S12.**
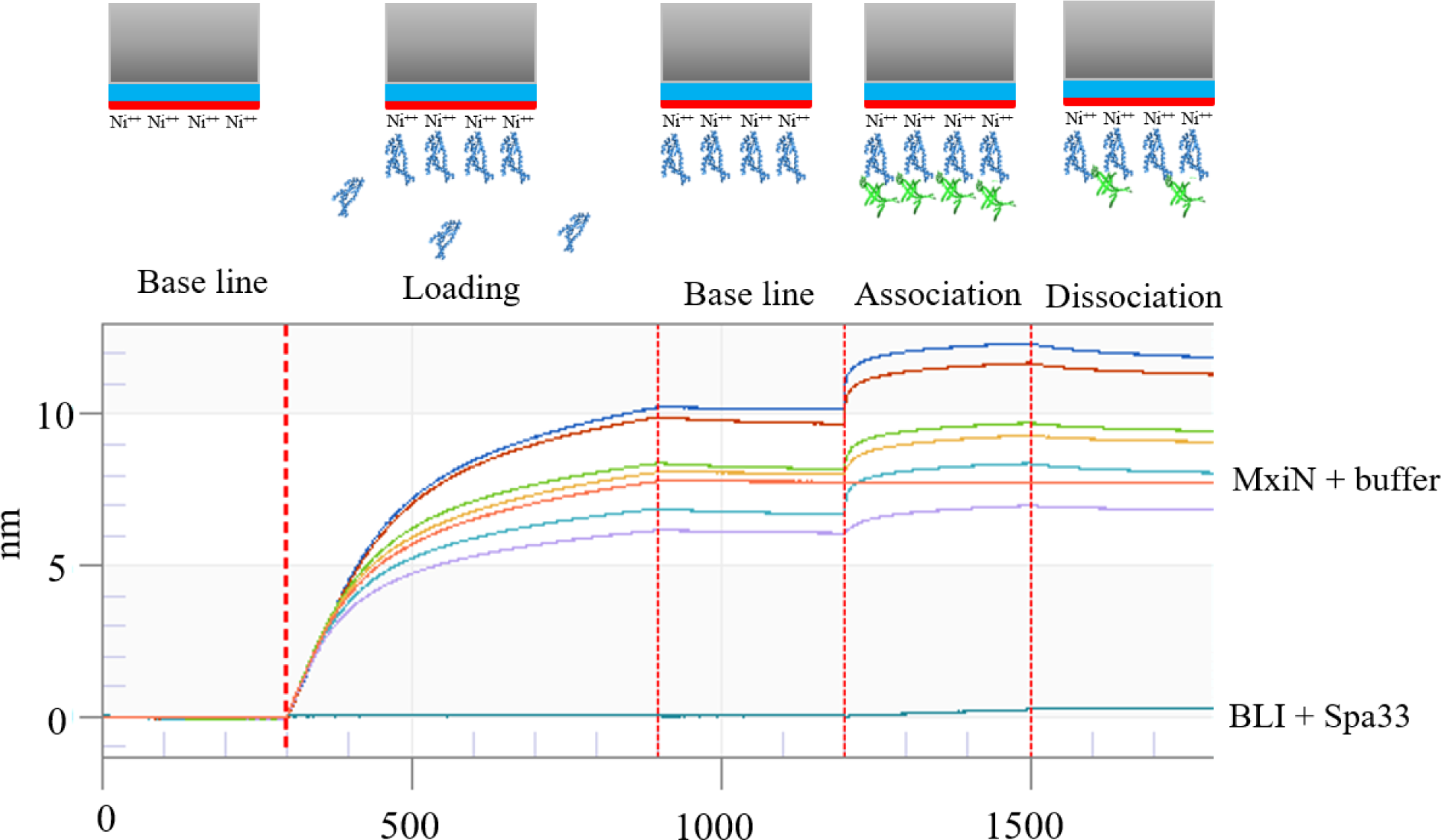
Biolayer Interferometry (BLI) analysis of Spa33 interacting with His_6_-tagged MxiN. MxiN with a His_6_-tag was loaded onto nickel-NTA biosensor tips and a baseline established. Association was then measured in buffer alone and with six different concentrations of wild-type Spa33 (heterotrimer) and the association monitored in real time. The MxiN-containing solution was then replaced with buffer alone and dissociation was then monitored in real time. The bottom baseline trace is the addition of Spa33 to sensor tips without prior loading with His_6_-tag MxiN.

**Supplemental Figure S13.**
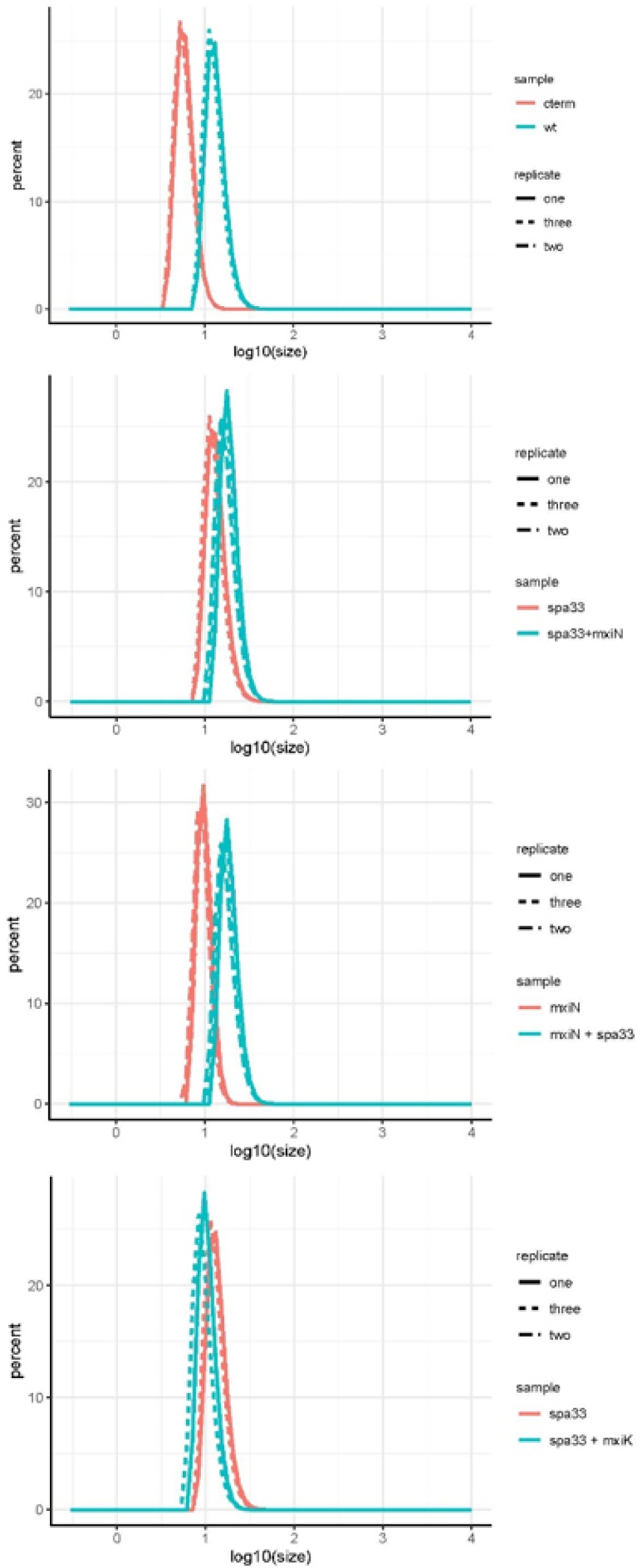
Dynamic light scattering of Spa33-containing complexes. Raw scans (in triplicate) are shown for wild-type Spa33 versus Spa33^C^ (**top panel**), Spa33 versus the Spa33-MxiN complex (**second panel**), MxiN versus the Spa33-MxiN complex (**third panel**) and Spa33 versus the Spa33-MxiK complex (**bottom panel**).

**Supplemental Figure S14.**
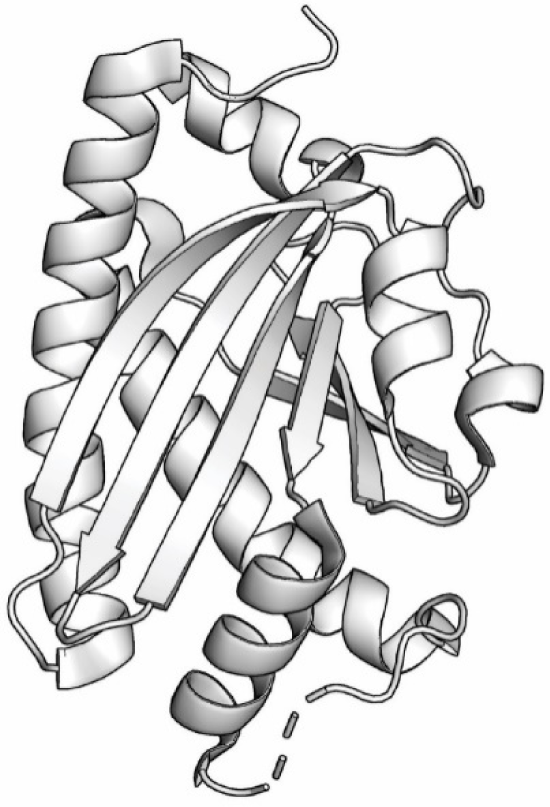
Crystal structure of FliM from *Thermatoga maritima*. The crystal structure *T. maritima* (PDB: 2HP7) has a greater α-helical content than does FliN and the CD spectrum of the wild-type Spa33 trimer suggests that this is also true for the N-terminal portion of Spa33^FL^.

**Supplemental Figure S15.**
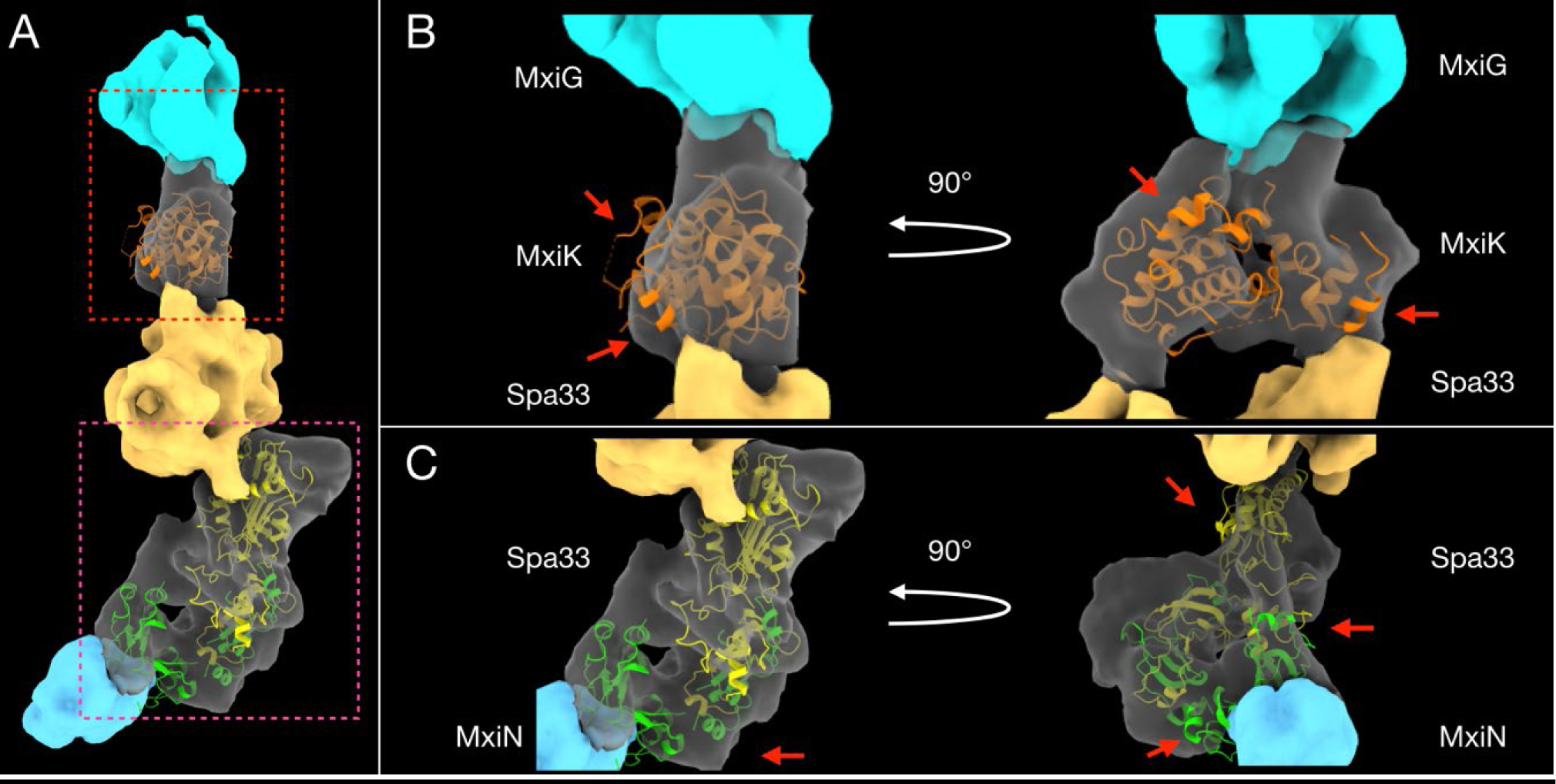
Up close model of an injectisome pod. **A)** The 3D surface rendering image of a single pod structure from **Panel B** of **Figure 7** is shown. **B)** The upper red square from **Panel A** is shown up close so that a fit of the crystal structure of *Pseudomonas aeruginosa* SctK can be more easily viewed. The sequence of SctK from *P. aeruginosa* is approximately 15-20% identical to MxiK based on analysis by Clustal Omega (Sievers et al. 2011. Mol. Syst. Biol. 7:539; Needleman and Wunsch 2970. J. Mol. Biol. 48:443-453) and most of its structure (as a monomer) readily fits into the rendering (Muthuramalingam et al. 2020. J. Mol. Biol. 432:166693. doi: 10.1016/j.jmb.2020.10.027). There are, however, a few areas (indicated by red arrows) that are located outside of the MxiK density. The **right side** is rotated 90°. **C)** The lower magenta square from **Panel A** is shown up close in **Panel C**. The model of the Spa33 heterotrimer, based on a model of the FliM/FliN heterotetramer, fits well within the lower of the two main pod densities.

